# Aging disrupts blood-brain and blood-spinal cord barrier homeostasis, but does not increase paracellular permeability

**DOI:** 10.1101/2024.02.12.580035

**Authors:** Mitchell J Cummins, Ethan T Cresswell, Doug W Smith

**Affiliations:** Neurobiology of Aging and Dementia Laboratory, School of Biomedical Sciences and Pharmacy, College of Health, Medicine and Wellbeing, University of Newcastle, Callaghan, NSW, Australia; Brain Neuromodulation Research Program, Hunter Medical Research Institute, New Lambton, NSW, Australia

**Author notes:** Corresponding Author: Doug Smith Address: University of Newcastle, Callaghan, NSW, Australia.

**Keywords:** Aging, blood-CNS barriers, Permeability, RNA-Seq, laser-capture, scRNA-Seq meta-analysis

## Abstract

Blood-CNS barriers protect the CNS from circulating immune cells and damaging molecules. It is thought barrier integrity becomes disrupted with aging, contributing to impaired CNS function. Using genome-wide and targeted molecular approaches, we found aging affected expression of predominantly immune invasion and pericyte-related genes in most CNS regions investigated, especially after middle age, with spinal cord being most impacted. We did not find significant perturbation of tight junction genes, nor were vascular density or pericyte coverage affected by aging. We evaluated barrier paracellular permeability using small molecular weight tracers, serum protein extravasation, CNS water content, and iron labelling measures. We found no evidence for age-related increased barrier permeability in any of these tests. We conclude that blood-brain (BBB) and blood-spinal cord barrier (BSCB) paracellular permeability does not increase with normal aging in mouse. Whilst expression changes were not associated with increased permeability, they may represent an age-related primed state whereby additional insults cause increased leakiness.

## 4 Introduction

Aging is associated with declining CNS function and compromised blood-CNS barrier integrity is thought to contribute to this decline (Bors et al., 2018; Goodall et al., 2018; Montagne et al., 2015). Aging is proposed to impair endothelial cell (EC) tight junctions and/or pericyte cell (PC) signalling, which are critical to barrier ‘tightness’, resulting in neurotoxic and neuroinflammatory factors gaining access to CNS parenchyma.

Two of the blood-CNS barriers, the blood-brain barrier (BBB) and the blood-spinal cord barrier (BSCB) are formed by vascular ECs, PCs, astrocyte end-feet, and secreted basement membrane, with associated intercellular junctions and signalling, (Abbott, Patabendige, Dolman, Yusof, & Begley, 2010; Daneman & Prat, 2015). Tight junctions between ECs limit the ability of molecules and immune cells to move between the ECs via the paracellular pathway. The transcellular pathway, on the other hand, requires passage through ECs, conferring regulated entry of needed and active or passive exclusion of unwanted molecules (Abbott et al., 2010; Daneman & Prat, 2015). Endothelial barrier permeability is influenced by PC signalling via the basement membrane, and astrocytic end-feet that surround CNS vessels.

Whilst blood-CNS barrier permeability has been extensively studied, a consensus has not been established as to whether permeability increases with aging. Human studies using the ratio of cerebrospinal fluid (CSF) to serum albumin as a measure of barrier permeability, generally report increases with age (Garton, Keir, Lakshmi, & Thompson, 1991; Pakulski, Drobnik, & Millo, 2000). However, a major limitation of this approach is lack of barrier specificity, as the measure is more of an indicator of blood-CSF barrier (BCSFB) and not BBB or BSCB function (Chen, 2011). Human studies using high resolution, contrast-enhanced MRI, have yielded equivocal results (Montagne et al., 2015; Montagne et al., 2020; Senatorov et al., 2019). Human post-mortem histological studies have found age-related increases in extravasated blood proteins in brain, which has been interpreted as indicative of BBB leakiness (Goodall et al., 2018; Senatorov et al., 2019).

In animal models, evidence for age-related increases in CNS barrier permeability is similarly mixed. Studies using exogenous tracers have reported no effects (Goodall et al., 2018; Ritzel et al., 2015), or an increase in tracer extravasation with aging (Hafezi-Moghadam, Thomas, & Wagner, 2007; Park et al., 2018; Senatorov et al., 2019). Endogenous blood protein extravasation has also been used in animals. Again, studies report either no effects (Bell et al., 2010), or increases in leakiness to blood proteins with aging (Elahy et al., 2015; Park et al., 2018; Senatorov et al., 2019; Soto et al., 2015; Yang et al., 2020). Another indicator of declining CNS barrier integrity is loss of pericyte coverage. Some studies have found up to 50% decline in pericyte coverage (Soto et al., 2015; Yang et al., 2020) while others have found no age effect (Bell et al., 2010; Park et al., 2018).

The reasons for these discordant findings are not obvious. To comprehensively understand the impact(s) of aging on blood-CNS barriers, we first carried out deep RNA-seq on cortex (CTX), and spinal cord (SC) from young and old age mice. This discovery-based approach was followed by more targeted analyses including additional CNS regions (hippocampus (HIP) and cerebellum (CB)), using an array of approaches spanning molecular through to functional, with a focus on assessing the paracellular pathway and endothelial cell tight junctions.

## 5 Results

### 5.1 Expression of immune invasion, pericyte, and extracellular matrix genes is perturbed by aging, but endothelial tight junction gene expression remains stable

#### 5.1.1 Aging blood-CNS-barrier transcriptomes indicate BBB/BSCB dysfunction

We deep sequenced RNA from CTX and SC of young and old mouse CNS. The sample read depth for the 100 bp, paired-end reads was 81.9 ± 2.3 (average ± SEM) million. The complete gene expression datasets for both CNS regions and DEGs from aging comparisons (q<0.05) and Gene Ontology (GO) enrichment analyses are in Supplementary File 1.

There were 173 enriched GOs in CTX and 761 in SC (enrichment score >2, FDR<0.01). Of these, 22 CTX and 36 SC enriched GOs were related to blood vessels or cell junctions (Supplementary Table 1), suggesting components of the BBB/BSCB are affected by aging. Fig. 1a shows a subset of barrier-related enriched GOs (>3-fold enrichment). To gain a more detailed understanding of which CNS barrier components may be affected by aging, DEGs were compared to experimentally-determined gene sets. We used barrier cell-type-enriched gene sets derived from improved cell-type-specific approaches and meta-analyses. To reduce variability in cell-type-specific gene sets we used an ‘at least in two gene sets’ criterion for gene inclusion (see Methods). This compilation approach yielded 360 endothelial-, 114 pericyte-, and 286 astrocyte-cell-specific genes for enrichment analyses (cell-specific-gene sets can be found in Supplementary Table 2). Fig. 1b shows the results of cell-type enrichment analyses, which indicate all barrier cell types are impacted by aging.

**Fig. 1:**
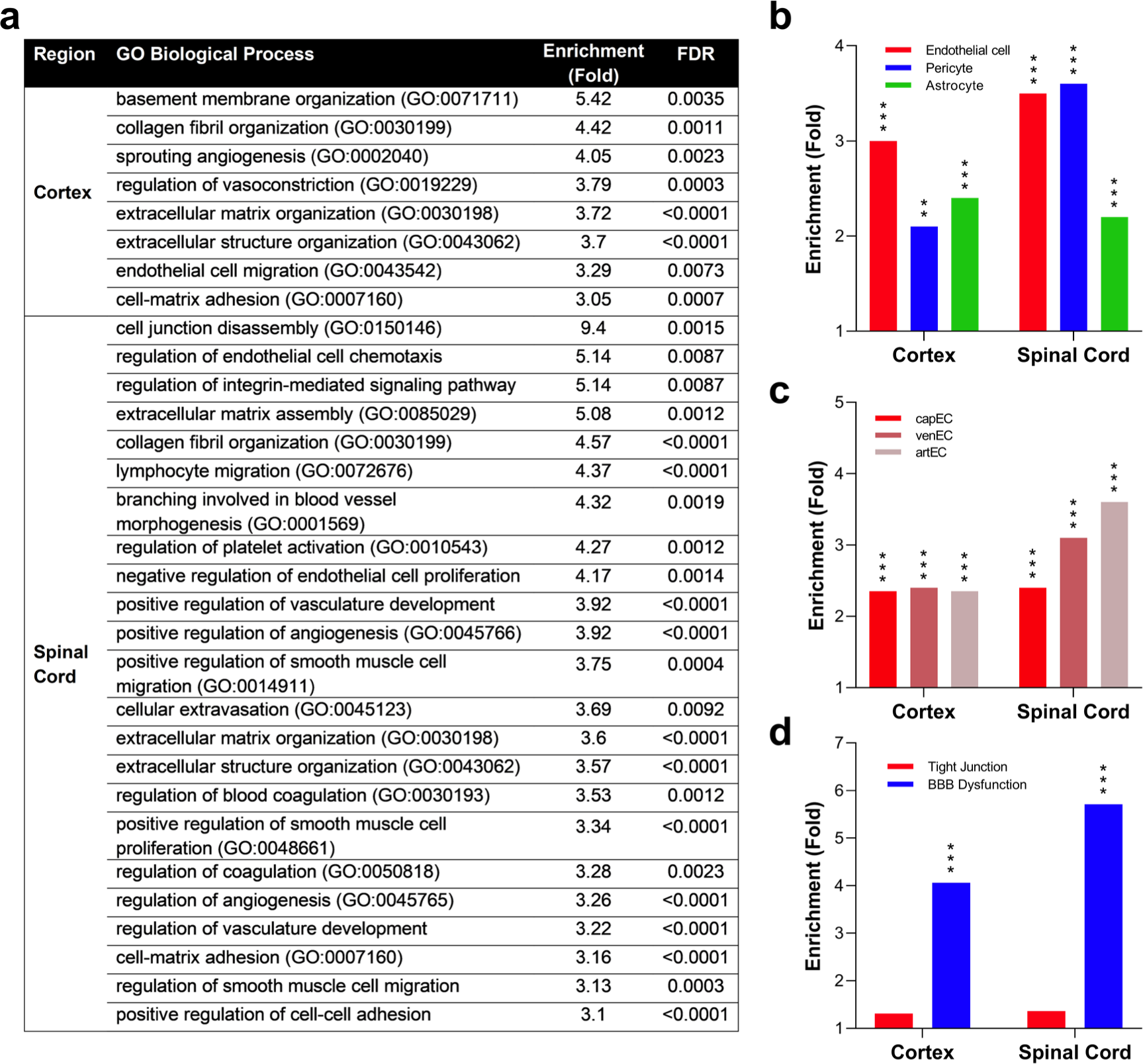
RNAseq of CTX and SC indicates barrier transcriptomes are perturbed by aging. **(a)** Barrier-related GO enriched in CTX and SC DEG sets with aging. GO with enrichment >3 and FDR < 0.01 are shown. **(b)** Enrichments plotted for endothelial- (EC), pericyte-, and astrocyte-cell-type-specific genes in the DEG sets for CTX and SC, using cell-type-specific gene sets as described in the text. **(c)** Enrichments plotted for ECs located at different parts of the cerebrovasculature tree. **(d)** Enrichments plotted for EC tight junction genes and BBB dysfunction module genes. See Methods for details on enrichment analyses. (**b-d**) **p<0.01 ***p<0.001.

Additionally, we made use of recently characterised cerebrovasculature tree-region-dependent EC molecular profiles (Vanlandewijck et al., 2018) to determine if aging impacts all regions of the tree similarly (Fig. 1c, Supplementary Table 3). Interestingly, arteriolar and venule ECs appear more impacted by aging in the SC compared to capillary ECs. This is intriguing as different tree regions are responsible for separate, albeit related, functions. In general, arterioles control blood flow and pressure, capillaries mediate entry and exit of molecules from the CNS parenchyma and modulate local blood flow in response to stimuli, and venules mediate immune cell invasion (Ashby & Mack, 2021; Daneman & Prat, 2015; Hall et al., 2014; Zlokovic, 2008).

Tight junctions between ECs are critical to barrier integrity. We used a 70-gene tight junction gene set (Munji et al., 2019), to assess impact of aging on EC tight junctions (Supplementary Table 4). There were no significant tight junction gene set enrichments in CTX or SC, indicating no aging effects (Fig. 1d, p>0.05). Munji et al. (2019) established a barrier dysfunction gene set comprising 136 genes upregulated in disease or injury (Supplementary Table 5). We found significant enrichment for BBB dysfunction genes (Fig. 1d), with 22% and 36% being differentially expressed in CTX and SC, respectively. All hypergeometric test results are in Supplementary Table 6. Overall, our RNA-seq analyses indicate blood-CNS barriers become perturbed with age, with SC being impacted to a greater extent than CTX. While all barrier cell types are affected, EC tight junction genes are not significantly altered.

#### 5.1.2 Barrier expression changes are most prevalent in the aging SC and occur post-middle age

To gain a more complete understanding of regional heterogeneity in barrier aging, we investigated age-related gene expression changes in 41 barrier-related genes in the CTX, corpus callosum (CC), hippocampus (HIP), cerebellum (CB), spinal cord grey matter (SCGM) and spinal cord white matter (SCWM), from the same animals using qPCR. Genes were functionally grouped, as follows: ECs (7 genes), PCs (10), immune cell invasion (6), extracellular matrix (ECM) (4), integrin-laminin binding (6), and genes involved in signalling between barrier cell types (8). Gene level expression results are presented in Fig. 2a. Supplementary Table 7 contains group fold changes, and p-values for all comparisons.

**Fig. 2:**
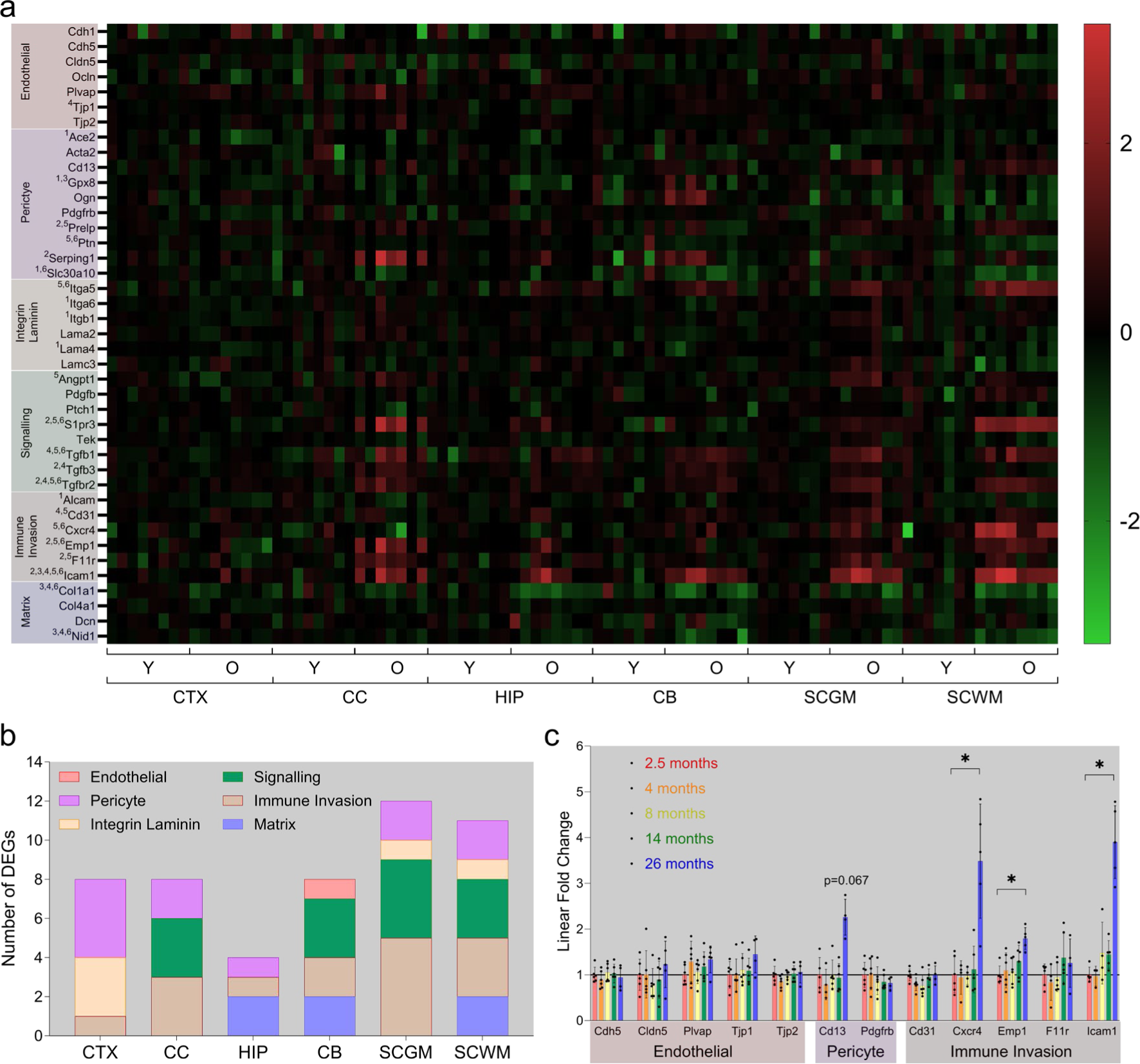
BSCB is more impacted by aging than BBB. **(a)** Heatmap of barrier gene expression. Individual sample expression is relative to average expression of young in matching CNS region. Scale bar is log_2_ FC. Superscripted numbers with gene names indicate region(s) reaching statistical significance: CTX^1^ CC^2^ HIP^3^ CB^4^ SCGM^5^ SCWM^6^. **(b)** Number of significantly differentially expressed genes (DEGs) of each functional group in each CNS region. **(c)** Relative expression of selected barrier genes across mouse adult lifespan (2.5, 4, 8, 14, 26 months) in SC. Expression is relative to 2.5-month group. Dots indicate individual animals. Bars indicate mean. Error bars ±SD. *p<0.05 corrected. CTX – cortex; CC – corpus callosum; HIP – hippocampus; CB – cerebellum; SC – spinal cord; GM – grey matter; WM – white matter.

Based on number of significant DEGs, the SC was the most affected region, with differential expression of 12/11 (GM/WM) of 41 genes. CTX, CC, and CB, were the next most impacted, with expression changes in 8 genes each. Only 4 genes were affected in HIP, a region generally considered to be of greater susceptibility to aging processes. No single gene was differentially expressed in all 6 regions, although immune signalling Icam1 was a DEG in 5/6 regions. Aging affected genes from all six functional groups, but in a region-specific way.

Immune cell invasion was the only functional group perturbed in all 6 regions, although DEGs in this group were not strictly the same. PC genes were the next most widely affected, with differential expression in all regions except CB. Of the signalling genes investigated, the Tgf-related pathway was most impacted, with the receptor Tgfbr2 being upregulated in 4 regions. Fig. 2b. summarises the impact of aging on functional groups across CNS regions investigated.

We also assessed the age-progression of barrier expression changes. As SC was the most aging-affected CNS region, we investigated BSCB expression changes across the mouse adult lifespan. Only 3 of the 12 genes assessed were differentially expressed. All were immune cell invasion related (Cxcr4, Emp1, Icam1), overexpressed, and significance was not reached until after 14 months (Fig. 2c). The apparent increase in Cd13 expression did not survive correction for multiple comparisons. The complete dataset is available in Supplementary Table 8. These results indicate impacts of aging become evident post-midlife.

#### 5.1.3. EC-PC-specific gene expression analysis through microdissection

To obtain a more direct measure of barrier gene expression, we extracted RNA from microdissected microvessels for qPCR analysis (Fig. 3). CNS blood vessels were immunolabelled for Col-IV, a collagen secreted by both EC and PCs. Microdissection of Col-IV-positive profiles therefore samples these two cell types. Fig. 3c-d show examples of pre- and post-laser microdissection of immunolabelled Col-IV profiles and demonstrate the high spatial and cellular resolution with this approach. Tight junction-related genes Cdh5, Cldn5, Ctnna1, Ctnnb1 and Tjp1 were not differentially expressed in CTX or SC GM microvessels (Fig. 3a-b), but the tight junction and immune related F11r (junction adhesion molecule A, Jam-A) decreased in SC GM microvessels only (Fig. 3e). The complete microvessel-targeted gene expression dataset is available in Supplementary Table 9.

**Fig. 3:**
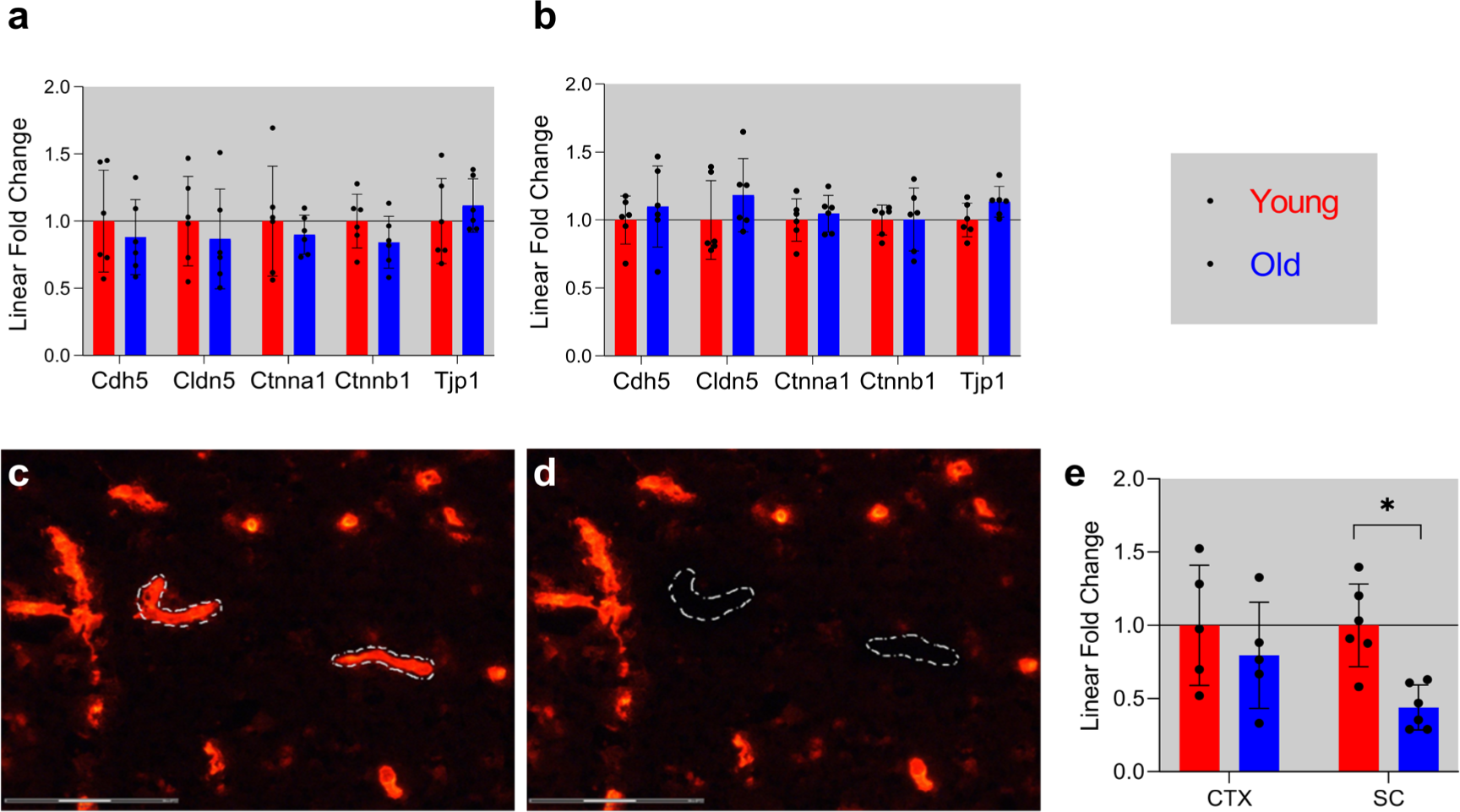
Laser microdissection of CNS microvessels. Relative expression of barrier tight junction genes in **(a)** CTX and **(b)** SCGM laser microdissected blood vessels. Col-IV-immunolabelled 10µm SC section **(c)** before and **(d)** after laser microdissection of microvessels, 40x objective. Scale bars are 75 µm. **(e)** Relative expression of F11r in CTX and SC blood vessels. Bars represent group fold change relative to young, points represent individual animals relative to young group average, error bars are SD for all graphs. n=5-6/group.* one-tailed Wilcoxon Mann-Whiney U Exact test corrected p-value < 0.05. CTX – cortex; SCGM – spinal cord grey matter.

#### 5.1.4 Meta-analysis of single-cell-RNA-seq studies

We complemented our molecular analyses with a meta-analysis of mouse whole brain single-cell datasets from five studies (Supplementary Table 10). Combining data from 152,696 cells improves statistical power and to date this is the largest number of single brain cells to be analysed for mouse brain aging. Gene expression data from 80,929 and 71,767 brain cells from young and old mice, respectively, were analysed using Seurat (Satija, Farrell, Gennert, Schier, & Regev, 2015). Twenty-one cell clusters were identified (Supplementary Table 11), with 2 overlapping clusters (0 and 7) of ECs (Fig. 4a-b), and two clusters (9 and 13) of vascular mural cells. Two main subclasses of mural cell populations, PCs and smooth muscle cells (SMCs), were also reported in a recent analysis of the murine brain cerebrovascular tree (Vanlandewijck et al., 2018). As PCs are comparatively rare CNS cells, past differential gene expression comparisons may have been underpowered. The present PC meta-analysis reduces this concern. The DEGs within each cluster are listed in Supplementary File 2.

**Fig. 4:**
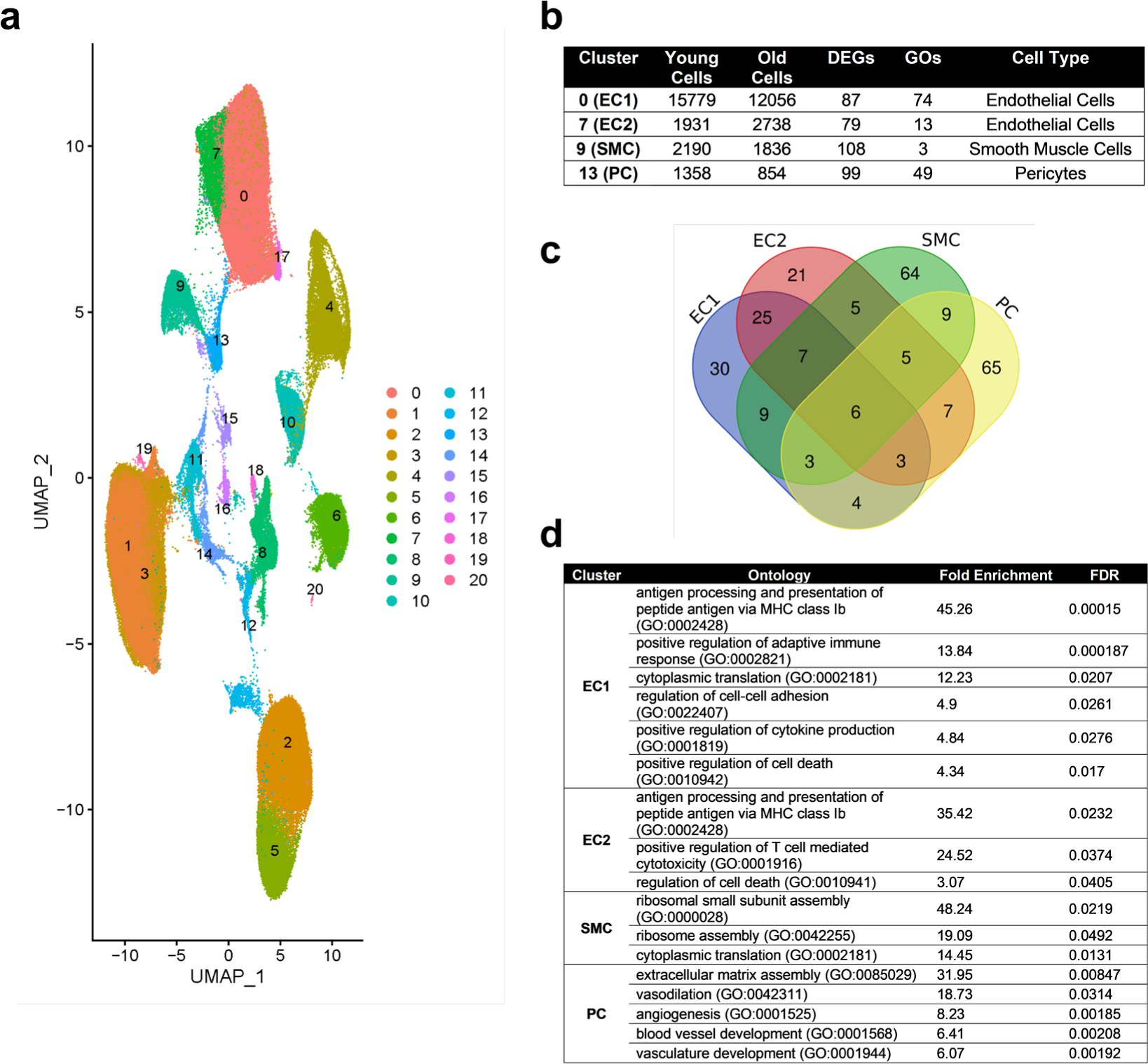
Aging brain single cell meta-analysis. **(a)** UMAP of aging brain cells. Putative cell type clusters: 0 - ECs; 1 - microglia; 2 - astrocytes; 3 - microglia; 4 - oligodendrocytes; 5 - astrocytes; 6 - oligodendrocyte precursor cells; 7 - ECs; 8 - neurons; 9 - smooth muscle cells; 10 - choroid plexus epithelial cells; 11 - perivascular macrophages; 12 - ependymal cells; 13 - PC; 14 - perivascular macrophages; 15 - vascular leptomeningeal cells; 16 - vascular leptomeningeal cells (ABC); 17 - olfactory ensheathing cells; 18 - neurons; 19 - proliferating; 20 - oligodendrocytes. **(b)** Number of cells in EC and mural cell clusters. **(c)** Venn diagram of EC and mural cell clusters age-related DEGs **(d)** Selected enriched GOs of DEGs in cluster 0 (EC1), cluster 7 (EC2), 9 (SMC), and 13 (PC). EC – endothelial cell; SMC – smooth muscle cell; PC – pericyte.

Mural cell (PCs and SMCs) clusters contained a greater number of DEGs (207) than EC clusters (166). Venn diagram analysis (Fig. 4c) showed little DEG overlap between PCs and SMCs (23 genes), whereas there was substantial overlap between the two EC clusters (41 genes). Relatively few DEGs overlapped different cell types and only 6 DEGs were common to all four cell clusters.

The EC clusters were screened for expression changes in genes that impact barrier function: the adherens (VE-cad, Pecam1, Nectin2, Afdn), tight (Tjp1-3, Ocln, Cldn5), and gap (Gja-e) junctions. Only Gja1 (a.k.a connexin 43) expression was affected, being lower with aging (Supplementary File 2), consistent with recent work (Zhan et al., 2023). Septin 2 (Sept2) is thought to organise EC junctional proteins (Kim & Cooper, 2021) and was upregulated in both EC clusters (Supplementary File 2). Interestingly, expression of junctional cadherin 5 associated (Jcad), a risk allele for coronary artery disease, was significantly increased in EC cluster 7. Jcad encodes a protein that disrupts EC function and promotes atherosclerosis (Douglas et al., 2020; Xu et al., 2019).

For mural cells we focussed primarily on mural cell-EC signalling that is critical for barrier formation and maintenance (Sweeney, Zhao, Montagne, Nelson, & Zlokovic, 2019). There were no significant expression changes in Pdgfb, Tgfb, Angpt/Tek, nor Ncad signalling genes (Supplementary File 2). However, there were a number of basement membrane genes that were age-affected including Col1a2, Col4a2, Lamb1, Spock2, Tnxb, Dmp1, Vtn, Nupr1, and Ogn. The collagens, Lamb1, Tnxb and Dmp1 had reduced expression while Spock2, Vtn, Nupr1, and Ogn were increased with aging.

GO analyses revealed that EC and SMC clusters were enriched for a small variety of GOs. Example enriched GOs are shown in the table in Fig. 4d for all clusters. Many EC cluster enriched GOs involve immune functions (40 of 87 enriched GOs for both EC clusters, FDR p<0.05), for example antigen processing and presentation, and adaptive immune system regulation involving T cells. Other EC enriched GOs involved cell death, protein processes, cellular responses to various stimuli, wound healing, and haemostasis. Consistent with there being separate EC clusters, cluster 0 had a greater number of and more diverse enriched GOs than EC cluster 7. While mural cell clusters had more DEGs than EC clusters, the number of enriched GOs was smaller (52, FDR p<0.05), indicating fewer biological process were perturbed by aging. Indeed, SMCs only had 3 significantly enriched GOs, all involving protein translation. PCs, however, were enriched for a diverse array of GOs including many related to blood vessel development and maintenance (14 of 49), the extracellular matrix (3/49), development, proliferation, and differentiation processes (8/49), synaptic signalling by gas (5/49), response to oxygen levels (1/49), and other general metabolic and cellular processes (see Supplemental Table 12 for the complete GO enrichments).

### 5.2 Aging subtly affects CNS blood vessel density but not pericyte coverage

Blood vessel density was assessed using Col-IV immunolabelling to identify vessels. The expected lower blood vessel density in WM compared to GM was observed (Ventura-Antunes & Herculano-Houzel, 2022), with density in WM being ∼50% of that observed in GM regions (Fig. 5a-e). Effects of aging were subtle, and only seen in two regions, with a small decrease in density in the CC (p=0.041) and a small increase in the CB (p=0.027). CTX, HIP, SCGM, and SCWM all showed no significant effects of aging on blood vessel density.

**Fig. 5:**
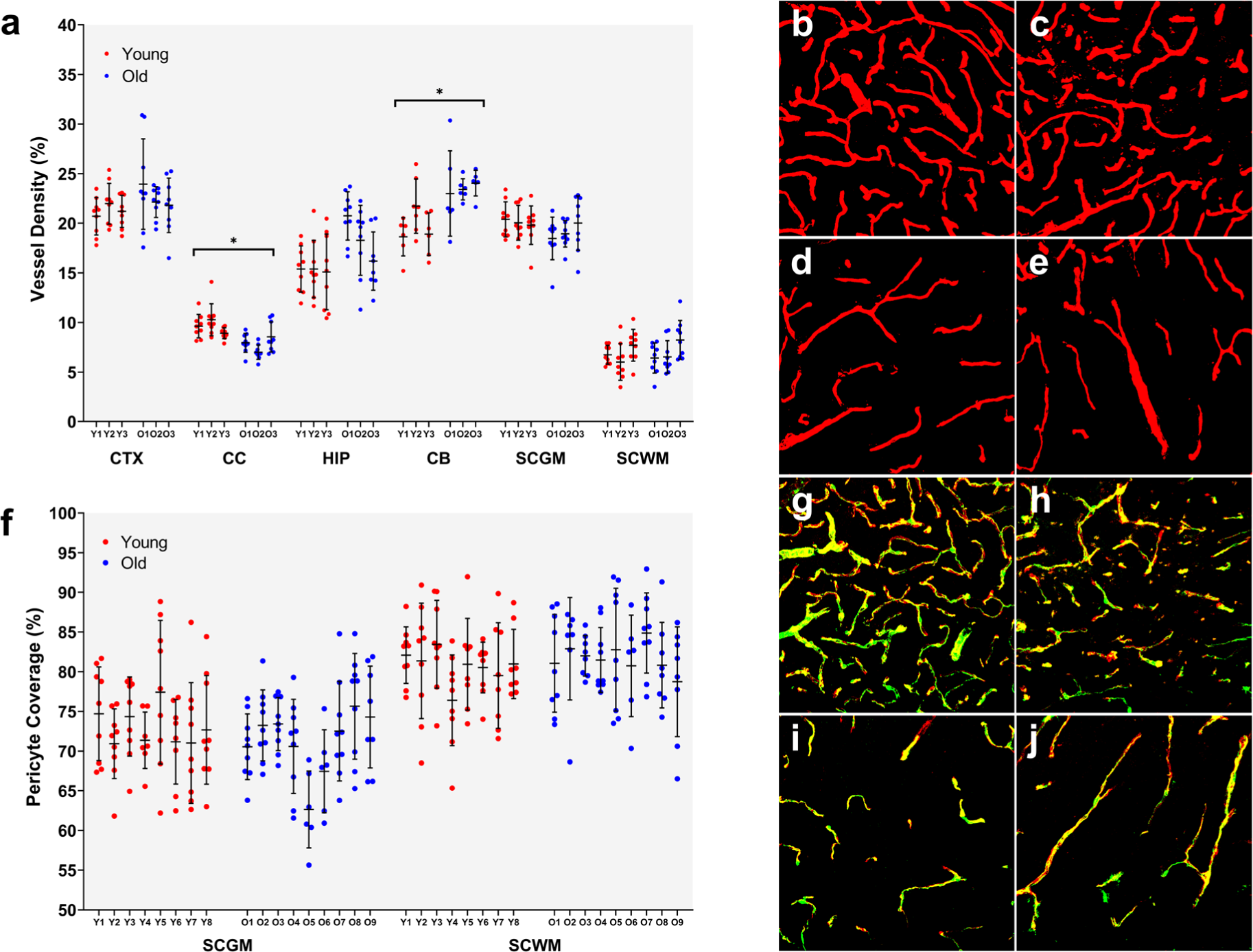
Blood vessel density and pericyte coverage in aging mouse CNS. **(a-e)** Blood vessel density. **(a)** Vessel density was quantified by Col-IV immunolabelling across 6 CNS regions. Points are individual measurements (6-9/animal). Error bars are mean ±SD. n=3/gp/region. * Nested t-test p < 0.05. **(b-e)** False colour processed images of Col-IV immunolabelling (red) in young **(b,d)** and old **(c,e)** spinal cord grey **(b,c)** and white **(d,e)** matter at 40x. **(f-j)** Pericyte coverage. **(f)** Pericyte coverage was quantified using Col-IV and CD13 double immunolabelling. Points are individual measurements (6-9/animal). Lines are mean and SD. n=8-9/gp/region. **(g-j)** False colour processed images of Col-IV (red) and CD13 (green) double (yellow) immunolabelling in young **(g,i)** and old **(h,j)** spinal cord grey **(g,h)** and white **(i,j)** matter at 40x. CTX – cortex; CC – corpus callosum; HIP – hippocampus; CB – cerebellum; SC – spinal cord; GM – grey matter; WM – white matter.

Pericyte coverage of blood vessels was assessed in the SC. Immunolabelling for CD13 was used to identify PCs and Col-IV immunolabelling to identify blood vessels (Fig. 5g-j). There were no significant differences between age groups in pericyte coverage in either the SCWM or SCGM (Fig. 5f). Pericyte coverage of blood vessels in WM appeared higher than in the GM, as expected (Winkler, Sengillo, Bell, Wang, & Zlokovic, 2012).

### 5.3 Functional analyses of blood-CNS barrier permeability reveal general lack of age effects

#### 5.3.1 Age-related loss of CNS water content

Brain and SC water content were determined using oven drying. Wet weight and dry weight both significantly increased with age in both regions (Supplementary Fig. 1a-b). Water content, expressed as a percentage of total (wet) weight, significantly decreased in the aged brain and SC (Fig. 6a).

**Fig. 6:**
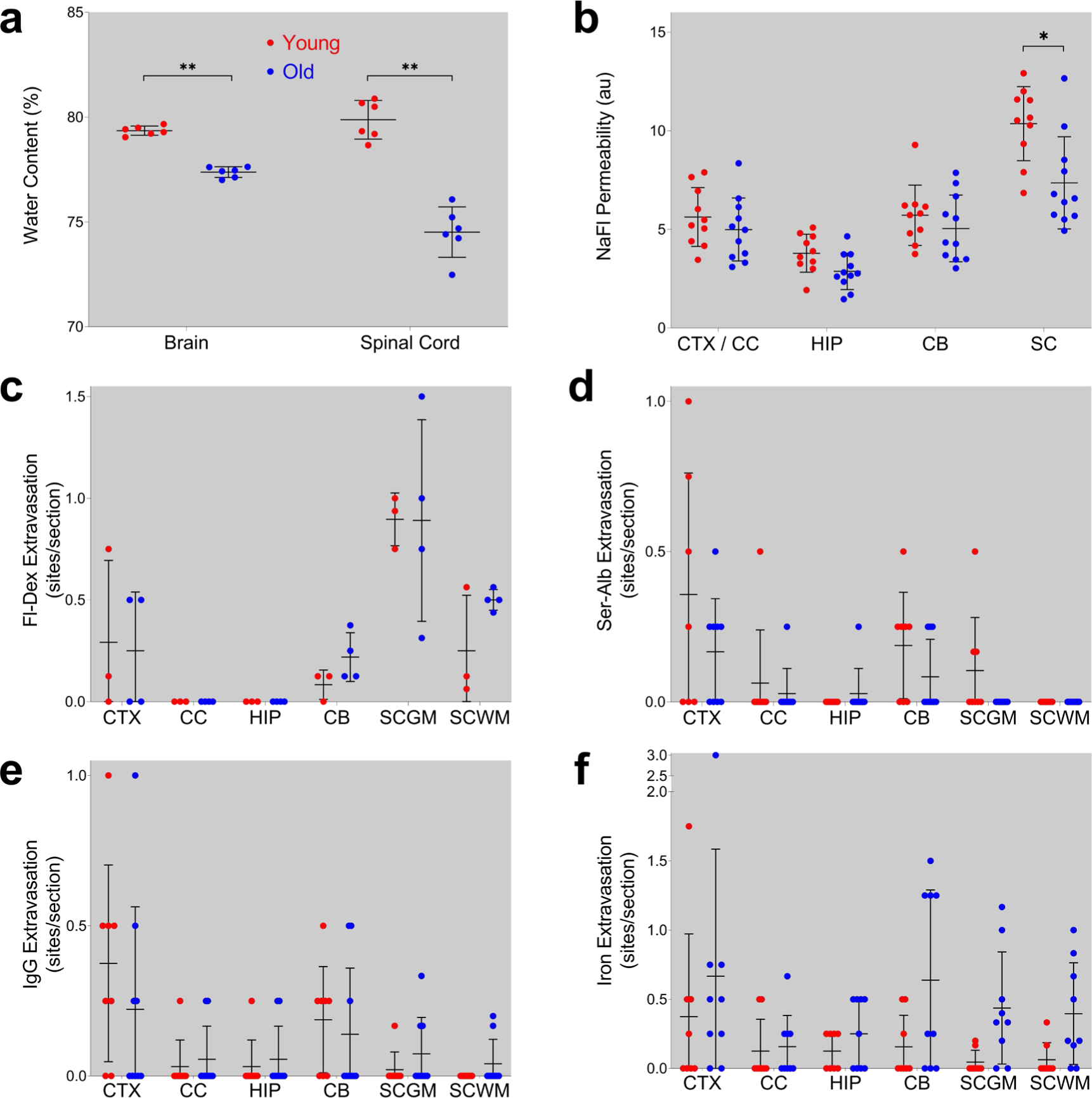
Functional assessment of the impact of aging on blood-CNS barrier permeability. **(a)** Effects of aging on brain and spinal cord water content expressed as percent of wet weight. n=6/gp. **(b)** Sodium Fluorescein (NaFl) permeability. n=10-11/gp. **(c)** Fluorescein dextran (Fl-Dex) permeability. n=3-4/gp. **(d)** Serum albumin (Ser-Alb) permeability. n=8-9/gp. **(e)** Immunoglobulin G (IgG) permeability. n=8-9/gp. **(f)** Iron labelling for microhaemorrhages. n=8-9/gp. Graphs: points represent individual animals; lines and error bars are mean ±SD; * one-tailed Wilcoxon Mann-Whitney U Exact test corrected p-value < 0.05. Legend in **(a)** holds for all graphs. CTX – cortex; CC – corpus callosum; HIP – hippocampus; CB – cerebellum; SC – spinal cord; GM – grey matter; WM – white matter.

#### 5.3.2 No age-related increase in blood-CNS barrier tracer permeability

BBB and BSCB permeability were measured using the low molecular weight tracer sodium fluorescein (NaFl, MW ∼376Da). CNS tissue and serum levels of NaFl were determined by fluorometry. Tissue levels were normalised to serum NaFl for each sample. Permeability did not change significantly in CTX, CC, or the CB, whereas it decreased in the SC (p=0.01) (Fig. 6b). The apparent decrease in the HIP did not survive correction for multiple comparisons (p=0.018 uncorrected, p=0.054 corrected).

Permeability was also measured using fluorescein dextran (MW ∼3kDa), a molecular weight tracer in the size range of small proteins. The number of areas demonstrating leakage (extravascular dextran) per tissue section was counted for CTX, CC, HIP, cerebellum grey matter (CBGM), cerebellum white matter (CBWM), SCGM and SCWM. Leakage indications were rare (Fig. 6c, and Supplementary Fig. 2a-d for images) and there were no significant age-related differences in any CNS region investigated.

#### 5.3.3 No age-related increase in endogenous indicators of CNS barrier dysfunction

We also investigated barrier permeability by labelling for endogenous indicators of barrier dysfunction. Brain and SC sections were immunolabelled for serum albumin (MW ∼68kDA) and IgG (MW ∼150kDA), and the number of areas with perivascular serum albumin (Fig. 6d) or IgG (Fig. 6e) per tissue section were counted. In both cases, perivascular labelling was rare (as for dextran experiments), and no regions had significant age-related effects (see Supplementary Figs. 3a-d and 4a-d for albumin and IgG images, respectively).

We also assessed barrier permeability by staining brain and SC sections using Prussian Blue to label iron, a marker of microhaemorrhages. Putative microhaemorrhage events were rare, as for serum albumin and IgG perivascular labelling, and similarly there were no age-related effects in the regions investigated (Fig. 6f, and Supplementary Fig. 5a-d for images). We did find an age-related effect in the thalamus (Supplementary Fig. 6a-b) with iron detected in old animals, which has previously been reported (Wang 2020, Taylor 2020) and serves as a positive control for the labelling.

## 6 Discussion

Blood-CNS-barrier disruption is thought to contribute to age-related decline in CNS function. However, a consensus on the effects of normal, non-pathological, aging on barrier integrity is yet to be reached. To address this, we have carried out a comprehensive assessment of BBB and BSCB, using an array of molecular, including a meta-analysis of scRNA-seq studies, and functional analyses.

Transcriptomic profiling and enrichment analyses indicated the three cell types most directly influencing barrier function, ECs, PCs, and astrocytes, were affected by aging. To more directly assess barrier properties at the molecular level, we did an enrichment analysis with an EC tight junction gene set (Munji et al., 2019), but found no significant enrichment, indicating aging does not overtly affect barrier tight junctions in CTX or SC. However, when enrichment was analysed with an EC dysfunction gene set, derived from conditions causing barrier leakiness (Munji et al., 2019), there was significant enrichment for both CTX and SC. Note, only one gene is common to both EC tight junction and dysfunction gene sets (Munji et al., 2019). Overall, this suggests aging makes CNS ECs dysfunctional but not necessarily leaky. Indeed, our scRNA-seq meta-analysis is consistent with this notion as we found little evidence for EC tight junction changes, yet a number of GOs were significantly impacted by aging, in particular those involving the immune system.

qPCR analyses also revealed variable effects of aging across the CNS, with the SC being the most affected overall, and HIP the least affected. The limited impact on the HIP is somewhat surprising given its known susceptibility to aging. Studies have reported aging-related hippocampal atrophy, functional decline (Bettio, Rajendran, & Gil-Mohapel, 2017; Gordon, Blazey, Benzinger, & Head, 2013), and impaired hippocampal-dependent cognitive performance (O’Shea, Cohen, Porges, Nissim, & Woods, 2016). Our findings suggest the hippocampal BBB is not a contributing factor to aging-related cognitive decline, which is consistent with recent human work that showed no correlation between age and indices for PC dysfunction and hippocampal BBB leakiness in either normal or mildly cognitively impaired cohorts (Nation et al., 2019).

EC tight junction gene qPCR analysis found just one significant change with age, with Tjp1/ZO-1 expression increased only in CB. Other studies have reported age-related decreases (Elahy et al., 2015; Mooradian, Haas, & Chehade, 2003; Murugesan, Demarest, Madri, & Pachter, 2012; Yang et al., 2020), or no change (Bell et al., 2010; Murugesan et al., 2012) in expression of tight junction genes/proteins. Our RNA-seq enrichment analyses and our scRNA-seq meta-analysis, support our qPCR finding of minimal tight junction changes. Overall, decreased CNS EC tight junction gene expression is not a widespread characteristic of normal aging.

For PCs, expression of the barrier-critical gene, Pdgfrb, was only significantly affected (decreased) in CTX. Expression of the heparin-binding, ECM, leucine-rich repeat protein (Prelp) was increased in CC and SC GM (and CTX p=0.06, SC WM p=0.05) of old mice. Blood-CNS-barrier function for Prelp is not known, but it does bind soluble complement inhibitor C4b-binding protein (Happonen, Furst, Saxne, Heinegard, & Blom, 2012), and may therefore be involved in the innate immunity complement response aspect of inflammaging. Interestingly, we found decreased pleiotrophin (Ptn) expression in SC GM and WM. Ptn is a PC-secreted neurotrophic factor that when administered exogenously to mice prevents the neuronal loss associated with PC degeneration (Nikolakopoulou et al., 2019). While Ptn loss alone is not thought to lead to vascular disturbances or neuronal loss in young healthy animals (Krellman, Ruiz, Marciano, Mondrow, & Croll, 2014; Nikolakopoulou et al., 2019), it may be more deleterious in the context of the inflamed, aging CNS.

Eight EC – PC signalling genes were also investigated, and while transforming growth factor beta (Tgfb) signalling was significantly age-affected, our laser microdissection data (Supplemental Table 9) and single-cell data (Supplemental File 2) suggest the Tgfb signalling is not microvessel derived.

Genes involved in immune cell invasion were generally increased in expression in old animals, with Icam-1 being the most markedly affected (up ∼5-fold) and most widely affected (5 regions). Icam-1 is expressed by ECs and is involved in leukocyte adhesion and transmigration into CNS parenchyma (Abadier et al., 2015). Notably, Icam-1 expression is increased in the inflamed CNS, suggesting peripheral immune cells may infiltrate. Based on molecular signatures, we (and others) have found the mouse CNS to be inflamed, however others have not observed markers suggesting an increased number of immune cells have infiltrated (Elahy et al., 2015; Ximerakis et al., 2019). Icam-1 is not the only molecule involved in leukocyte transmigration into the CNS and the increased expression we report here may be indicative of a priming event that requires additional inflammatory factors before immune cells can extravasate. Others have reported increased numbers of infiltrating immune cells in the aged CNS, but also reported increased permeability to an injected dextran and decreased gene expression of Cdh5, Ocln, Tjp1, and Cldn5 (Propson, Roy, Litvinchuk, Kohl, & Zheng, 2021). We did not see increased permeability to a smaller dextran, nor did we find robust decreased expression of tight junction genes.

In the last part of our molecular analyses, we determined the time course of changes across the adult mouse lifespan. We quantified expression of a subset of genes that represented multiple aspects of barrier cell types and function in the SC, the most affected region as determined in qPCR analyses, and an understudied CNS region in aging research. Of the 12 genes assessed, only three were significantly changed and were overexpressed at the oldest age only, with all three being immune cell invasion related (Cxcr4, Emp1, Icam-1). Overall, these age-course data indicate that age-related changes in the BSCB occur in the latter half of the adult life span.

In summary, our molecular data yielded limited support for perturbation of tight junctions between ECs, although aging did impact each of the cell types that comprise the blood-CNS barriers, possibly establishing a ‘primed’ state. Functional implications, as assessed by biological processes gene ontology enrichment, indicated expression of genes involved in signalling between barrier cells and immune cell invasion were the most widely and markedly affected, whilst the molecular effects appear to be occurring relatively late in adult life. Overall, the effects aging on gene expression were typically modest, so we then sought to determine if morphological and/or functional characteristics of the blood-CNS barriers were impacted.

Previous studies have suggested blood vessel density is reduced in the aged CNS (Murugesan et al., 2012; Park et al., 2018; Reed et al., 2017), although others have reported no change (Bell et al., 2010; Goodall et al., 2018; Yang et al., 2020). While we found the expected WM vessel density being approximately 50% of that in GM, there was little change in density with aging, the exception being a decrease in CC, and increase in CB (Fig. 5a-e). Thus, we do not observe the changes previously seen, where marked decreases in vessel density were reported (Park et al., 2018; Reed et al., 2017). Another factor in barrier function is PC coverage, and our molecular data indicates potential for PC dysfunction with aging. Consistent with previous reports, we did not find age-related loss of PC coverage (Bell et al., 2010; Bors et al., 2018; Goodall et al., 2018; Park et al., 2018). We did find PC coverage appeared greater in WM, consistent with previous reports (Winkler et al., 2012). There is inconsistency across studies with reports of decreased pericyte coverage with aging (Soto et al., 2015; Yang et al., 2020). Taken together, our vessel density and PC coverage data indicate aging does not have a substantial impact on microvessel gross structure, suggesting blood vessels are mostly intact. There are, of course, well established changes in microvessel ultrastructure that occur with aging (Ceafalan et al., 2019). With our molecular and morphometric analayses indicating relatively limited impact of aging on blood-CNS-barriers, we then carried out a number of complementary functional analyses.

Tissue water content is one measure of barrier permeability. Increased tissue water is indicative of barrier leakiness (Armulik et al., 2010). Contrary to expectation, we found significant decreases in water content in both the brain (down ∼2%) and spinal cord (down ∼5%). This is in stark contrast to data from Park et al. who reported a ∼4% increase in brain water content with age (Park et al., 2018). We found a number of differences between ours and the Park study. For example, they found an ∼50% decrease in cortical microvessel density, whereas we found no age-related changes. They also found increased extravascular NaFl whereas we did not. Our data are consistent with a previous report of decreased extracellular space with age (Sykova et al., 2002).

Exogenous tracers of different molecular weights were also used to assess barrier permeability. We found no effects of age with relatively small (NaFl) and large (dextran) molecular weight tracers, with the exception of a significant decrease in extravasated NaFl in SC of old animals. This decrease is likely not due to a decrease in vessel density as we did not find an age effect on density in SC. Consistent with previous reports, we did find the BSCB to be more permeable than the BBB (Ritzel et al., 2015; Winkler et al., 2012). Ritzel et al. (2015) also found a similar SC outcome, with lower NaFl permeability in aged mice (Ritzel et al., 2015). Park et al. (2018) found an ∼50% increase in brain NaFl permeability in old mice. Outcome differences between the Park and current studies are difficult to reconcile.

Exogenously administered barrier tracers may not recapitulate the physiology of endogenous blood-borne factors (Yang et al., 2020). Therefore, we also analysed the degree to which endogenous IgG, serum albumin, and hemoglobin iron (an indicator of hemorrhage) was located extravascularly across CNS regions. As with the exogenous tracers, there were no increases in extravascular labelling with aging. One exception was increased iron labelling in the thalamus of old mice, which is consistent with previous reports of increased microbleeds in the aged thalamus (Taylor et al., 2020; Wang et al., 2021). Others have reported increased extravascular labelling in aged mice, but only after induction of an inflammatory event. supporting the notion of a “primed”, but not “leaky”, state (Sumbria et al., 2018).

In summary, our functional findings support our molecular data, indicating the blood-CNS barriers investigated do not get leakier with aging. If anything, the BSCB becomes less permeable to small molecules, as indicated by our tracer results, and supported by decreased water content. Aging-related increases in basement membrane thickness (Ceafalan et al., 2019) could contribute to loss of permeability. Basement membrane is made up of extracellular matrix proteins secreted by the three component cell types of the barriers. Aging-related changes in basement membrane thickness and/or relative abundances of extracellular matrix proteins could affect barrier permeability.

Considering the present findings, along with those of the many previous studies investigating the effects of aging on blood-CNS-barriers, a critical issue is apparent, that being, the marked discrepancies between studies, even when the same strain and gender of inbred mice, and types of assays were used. For example, Park et al. (2018) used C57BL/6 mice as we have in the present study and they report vastly different effects of aging, including age-related increases in brain water content, extravascular endogenous and exogenous markers, and marked neurodegeneration and overall brain shrinkage (Park et al., 2018). None of which we found (Fig. 6 and Supplementary Fig. 7). There are many such conflicting examples in the literature and this inter-study variability makes it difficult to arrive at a consensus on the impact of aging on CNS barriers. It may be that other factors need to be considered and reported in these types of barrier studies. For example, diet is known to affect barrier function, with diets enriched in saturated fatty acids or cholesterol showing increased serum IgG extravasation (Takechi, Pallebage-Gamarallage, Lam, Giles, & Mamo, 2013). The gut micobiota can also affect blood-CNS-barrier permeability either directly with bacterial metabolites, or indirectly through peripheral immune cell activation (Kadowaki & Quintana, 2020; Wekerle, 2017). For example, it has been shown that short chain fatty acids produced by the gut microbiota impact the function of astrocytes and microglia (Erny et al., 2015), which could in turn alter CNS barrier permeability.

In conclusion, our data does not support a loss of barrier tight junctions in aging CNS, but they do indicate development of an inflammatory profile that may render the barriers susceptible to CNS and/or peripheral inflammatory events. In other words, normal aging may constitute a molecular ‘first hit’, with additional insults required for a barrier breach, at least by the paracellular pathway. Whatever the reasons for the discrepancies between ours and previous findings, there remains much to do before the effects of aging on the blood-CNS-barriers are fully understood. Indeed, recent work using a novel ‘endogenous tracer’ approach where blood proteins were labelled, demonstrated the transcellular pathway is impacted by aging (Yang et al., 2020), indicating the two pathways from the circulation into the CNS parenchyma maybe differentially impacted by aging.

## 7 Experimental Procedures

Detailed procedures can be found in Supplementary File 3.

### 7.1 Animals

Young (2-4 months) and old (>24 months) C57BL/6 male mice were used for all experiments. All animal work was undertaken in strict accordance with the University of Newcastle Animal Ethics Committee, and New South Wales and Australian animal research guidelines.

### 7.2 Tissue Dissection and Cryosectioning

For molecular analyses, mice were perfused with PBS, brains and spinal cords were frozen in isopentane on dry ice. Cryosections were collected onto microscope slides. All reagents and procedures were RNase free and carried out on ice where appropriate.

For immunolabelling and staining analyses, mice were perfused with PBS, followed by 4% paraformaldehyde (PFA). Tissues were post-fixed, cryosectioned at 40µm, and stored in PBS at 4°C.

### 7.3 Molecular Analyses – Gene expression

#### 7.3.1 Bulk RNA Sequencing

a) Tissue preparation for bulk RNA sequencing

RNA was extracted from cervical spinal cord and frontal cortex. DNA-free, RNA samples were sent to the Australian Genome Research Facility for sequencing.

b) Analysis of CNS barrier related gene expression by RNA sequencing

RNA sequencing was carried out to generate lists of differentially expressed genes (DEGs) for analyses for enrichment of CNS barrier-related genes. Total RNA was prepared for sequencing using Illumina platforms and reads were aligned using STAR. Aligned files were assessed to determine DEGs using Cuffdiff. DEG lists for the cortex and spinal cord were compiled using a FDR cut-off of <0.05. DEG lists were analysed for Gene Ontology (GO) enrichment using the PANTHER Overrepresentation Test for biological process (https://geneontology.org/). DEG lists were compared with the CNS barrier-related gene lists in Supplementary Tables 2-5. Enrichment of CNS barrier-related gene sets in the DEG lists was determined using a hypergeometric overlap calculator: https://systems.crump.ucla.edu/hypergeometric/. p-values were corrected for multiple comparisons.

#### 7.3.2 Age-related barrier gene expression across CNS regions by qPCR

All qPCR primer sequences for all experiments are available in Supplementary Table 13. Sample RNA integrity measured using the 3’ to 5’ ratio for all experiments is available in Supplementary Table 14.

To confirm RNA-seq outcomes and to add white matter (WM) regions to our analysis, qPCR analyses were carried out on grey (GM) and WM CNS regions. Brains and spinal cords were cryosectioned at 100µm thickness and cortex, corpus callosum, hippocampus, cerebellum, cervical spinal cord GM, cervical spinal cord WM, dissected out, RNA extracted and reverse transcribed.

Genes for qPCR analysis were selected based on importance to CNS barrier function, and on our RNA-seq results. qPCR reactions were run on Applied Biosystems 7500 or QuantStudio 6 Pro devices.

Relative expression differences between age groups were determined using the comparative Ct method. Statistical significance was determined using one-tailed Wilcoxon Mann-Whiney U Exact test. A p-value of p<0.05 was applied. All p-values were adjusted for multiple comparisons using the sequential Holm-Bonferroni procedure.

Based on RNA-seq and qPCR, the spinal cord was the most affected region and was the focus of time course analysis. Mice across five age groups (2.5 months, 4 months, 8 months, 14 months, and 26 months) were used. RNA was extracted and reverse transcribed from whole cervical spinal cords. qPCR reactions were run as described above. Differences between age groups were determined using the Kruskal-Wallis Test followed by a post-hoc Steel-Dwass Test with the 2.5 months old group as the control, with p<0.05 for significance.

#### 7.3.3 CNS blood vessel-specific gene expression using laser microdissection and qPCR

Brains and spinal cords were cryosectioned at 10µm. Sections were acetone fixed and incubated overnight in rabbit anti-collagen IV (ColIV) antibody in 2M NaCl PBS for RNA protection. After rinsing, sections were incubated with Alexa Fluor 594 conjugated donkey anti-rabbit antibody in 2M NaCl PBS. Immediately prior to microdissection, slides were dehydrated and delipidated. Microdissection was carried out using a PALM MicroBeam system (Zeiss). Blood vessel profiles were collected from GM regions only. qPCR and data analyses were done as above.

#### 7.3.4 Single cell meta-analysis

Publicly available gene-count data and metadata from all studies was downloaded from the Gene Expression Omnibus (see Supplementary Table 10 for series identifiers). Data was analyzed in R/4.0.1 using Seurat v3.2.2. Young mice were 2-3 months old and old mice were 18-24 months old.

For each study, cells with fewer than 200, and greater than 5,000 expressed detected genes were removed. The Seurat tutorial (https://satijalab.org/seurat/articles/integration_introduction.html) for dataset integration was followed. In brief, each dataset was normalized separately using NormalizeData, and features were selected using FindVariableFeatures “vst” selection method selecting 2,000 features. Features that were variable across datasets were selected using the SelectIntegrationFeatures. Anchors for integration were identified using FindIntegrationAnchors datasets were integrated using IntegrateData. Data was scaled and centered using ScaleData.

Clustering and visualization using the RunUMAP, FindNeighbours, and FindClusters was performed with defaults except for FindClusters with 0.3 resolution. Cluster marker genes were identified using FindAllMarkers for positive markers with minimum logfc.threshold of 0.25. Young and old cells in each cluster for each study were counted using the subset function to select age group and study name, and table function to select cell cluster metadata for the cell subset. Clusters were identified by searching for cluster marker genes using 2 databases (https://www.brainrnaseq.org/ and http://mousebrain.org). Differential expression testing between young and old cells in each cluster was performed using FindMarkers with grouping by “Age” and subset by cell cluster, with a minimum logfc.threshold of 0.25.

### 7.4 Blood vessel density and pericyte coverage

For blood vessel density, fixed sections from each CNS region were immunolabelled for ColIV. Sections were rinsed and incubated with AF594 fluorescent secondary antibody. Sections were washed, mounted in gelvatol, coverslipped, and imaged on a Nikon D-Eclipse C1 confocal microscope. Z-stacks were captured using a 40x objective, and the area of ColIV labelling relative to total area quantified using ImageJ. Differences between groups were assessed using a nested t-test (sample nested within the age group), with a p<0.05 for significance of the age effect.

Pericyte coverage was determined for SC by co-immunolabelling for CD13 and ColIV. Sections were confocal imaged as above. Relative areas covered by ColIV and CD13 were quantified using ImageJ as for vessel density. Percentage area of pericyte coverage was calculated using: ((area of CD13 labelling / area of ColIV labelling) * 100). Differences between groups were assessed using a nested t-test, with a p<0.05 for significance of the age effect.

### 7.5 Functional Analyses

Group differences for all functional analyses were assessed using a one-tailed Wilcoxon Mann-Whitney U Exact test, with a corrected p-value of <0.05.

#### 7.5.1 CNS Water Content

Brains and spinal cords were extracted, weighed to determine tissue wet weight, then dried at 85°C for 7 days. Wet weight, dry weight, and percentage CNS water content ((wet-dry)/wet*100) were calculated.

#### 7.5.2 Exogenous CNS Barrier Tracers

a) Sodium Fluorescein (MW 376.3Da)

Mice were injected i.p. with 200µl 6% NaFl and euthanized after 15 minutes. Blood was collected, animals perfused with PBS, and samples from frontal CTX, CC, HIP, CB, and cervical SC, were weighed, frozen on dry ice, and stored at -80°C. Serum was removed and stored at -80°C.

Tissue proteins were precipitated with tri-chloroacetic acid in preparation for NaFl fluorometry on a microplate reader. Sample fluorescence was normalised to sample serum fluorescence and sample tissue weight.

b) Fluorescein Dextran (MW 3kDa)

Mice were injected i.p. with 400µL 5mg/mL 3kDa lysine fixable fluorescein dextran and euthanized after 10 minutes. Tissues were drop-fixed overnight in 4% PFA. Brain and spinal cord sections were visualized on an Olympus BX51 epi-fluorescent microscope. The number of extravascular leakages was counted and normalised to the number of sections assessed.

#### 7.5.3. Endogenous Indicators of CNS Barrier Function

a). Serum Albumin (MW approx. 69kDa)

Brain and spinal cord sections were co-immunolabelled for mouse serum albumin and CD31. Sections were assessed on an Olympus BX51 epi-fluorescent microscope and the number of areas demonstrating perivascular serum albumin were counted and normalised to the number of sections counted.

b) IgG (MW approx. 150kDa)

Brain and spinal cord sections were prepared for co-immunolabelling for ColIV and IgG, and sections were assessed as described for serum albumin labelling.

c) Iron labelling for microhaemorrhages

Brain and spinal cord sections were stained for iron using potassium ferrocyanide. Dehydrated and delipidated sections were mounted in ultramount and assessed on an Olympus BX51 microscope under bright field illumination. The number of microhaemorrhages were counted and normalised as described for serum albumin labelling.

## Supporting information

Supplementary File 1 Bulk RNA-seq Results

Supplementary File 2 Single cell cluster aging-related DEGs

Supplementary File 3 Detailed experimental procedures

## 8.#Acknowledgements

We would like to acknowledge Dr Mark Bigland and Dr Gemma Parkinson for technical assistance with RNA sequencing sample preparation. We would like to acknowledge the assistance of Mr Aaron Scott for IT support in relation to RNA sequencing analyses. We thank the lab of Professor Liz Milward for the Potassium Ferrocyanide. We would like to acknowledge the Hunter Medical Research Institute, and the Greaves Family for providing project funding. We would like to acknowledge the University of Newcastle for funding supporting the aging mouse colony.

## 10. Author Contributions

MC was involved in the planning and performing of all experiments, analyses, and writing. DS was involved in the planning and performing of all experiments, analyses, and writing. EC reviewed and edited the manuscript, performed confocal microscopy, and was involved in RNA extractions and cDNA syntheses.

## 11. Data Availability

All data is available upon request to the authors. RNA sequencing reads are available in BioProject accession number PRJNA1073059.

## 13. Supporting Information Listing

Supplementary File 1 Bulk RNA-seq results.

Supplementary File 2 Single cell DEGs for all clusters.

Supplementary File 3 Detailed experimental procedures.

Supplementary Table 1 RNA-Seq GO enrichment.

Supplementary Table 2 Cell-type marker genes for DEG enrichment and hypergeometric test.

Supplementary Table 3 Cerebrovascular tree genes for DEG enrichment and hypergeometric test.

Supplementary Table 4 Tight junction genes for DEG enrichment and hypergeometric test.

Supplementary Table 5 Barrier dysfunction genes for DEG enrichment and hypergeometric test.

Supplementary Table 6 Hypergeometric test results.

Supplementary Table 7 Results for qPCR of barrier genes from multiple CNS regions.

Supplementary Table 8 Results for qPCR of barrier genes from spinal cord across lifespan.

Supplementary Table 9 Results for qPCR of barrier genes from laser microdissection blood vessels.

Supplementary Table 10 Single cell studies used for meta-analysis.

Supplementary Table 11 Single cell clusters.

Supplementary Table 12 Single cell GO enrichment for EC, PC, and SMC clusters.

Supplementary Table 13 qPCR primers.

Supplementary Table 14 3’:5’ sample ratios.

Supplementary Figure 1 Wet and dry weight of the aging brain and spinal cord.

Supplementary Figure 2 Dextran in the aging CNS.

Supplementary Figure 3 Serum albumin labelling in the aging CNS.

Supplementary Figure 4 IgG labelling in the aging CNS.

Supplementary Figure 5 Iron labelling in the aging CNS.

Supplementary Figure 6 Iron labelling in the aging thalamus.

Supplementary Figure 7 Gross measurements of the aging brain.

