## Supplementary Figures for "Aging disrupts blood-brain and blood-spinal cord barrier homeostasis, but does not increase paracellular permeability"

**Figure 1:** Wet and dry weight of the aging **(a)** brain and **(b)** spinal cord. Points are individual animals. ** p-value <0.01.

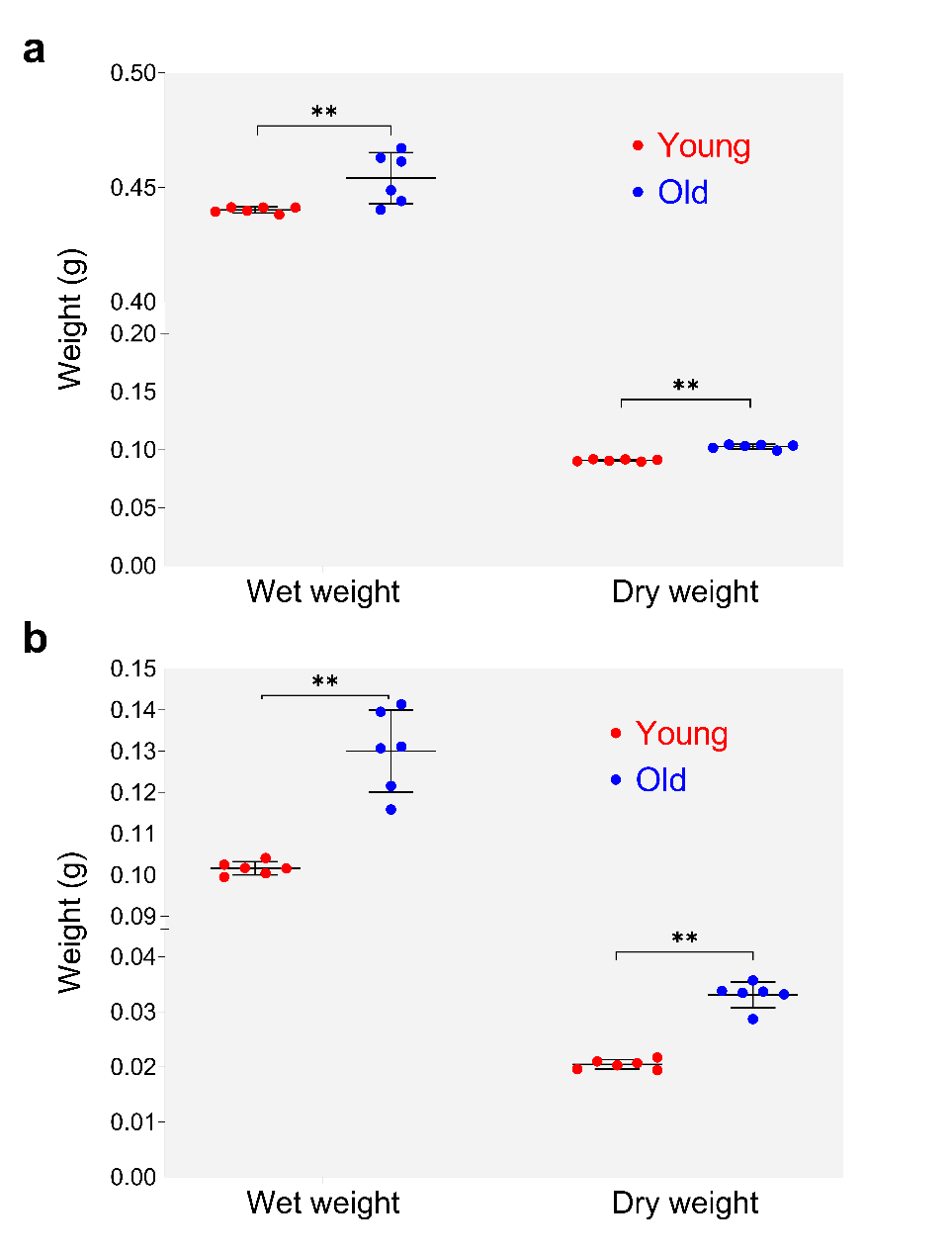

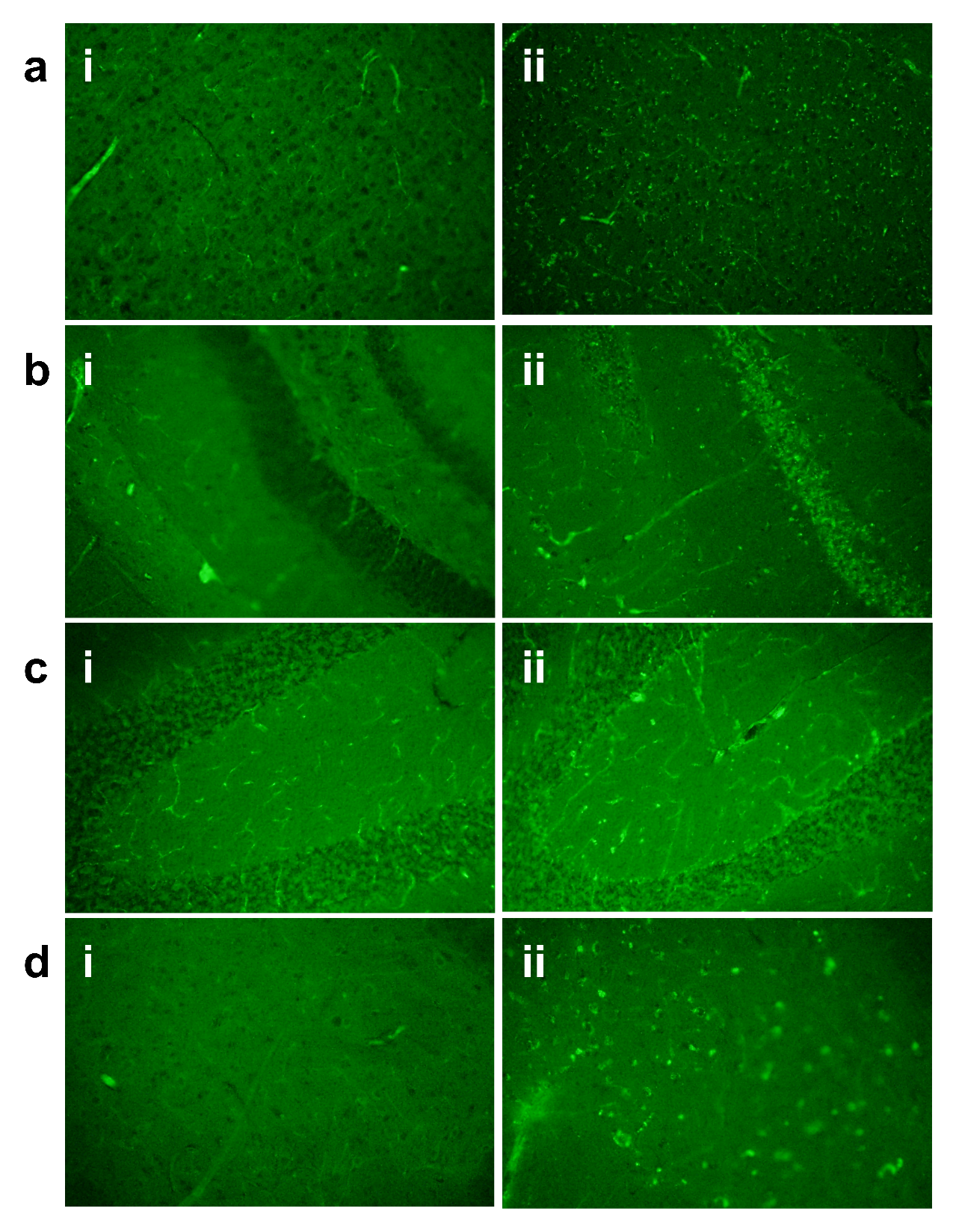

**Figure 2:** Dextran in the aging CNS. Example image of dextran distribution in the **(a)** cortex, **(b)** hippocampus, **(c)** cerebellum, **(d)** spinal cord in **(i)** young and **(ii)** old. 20x.

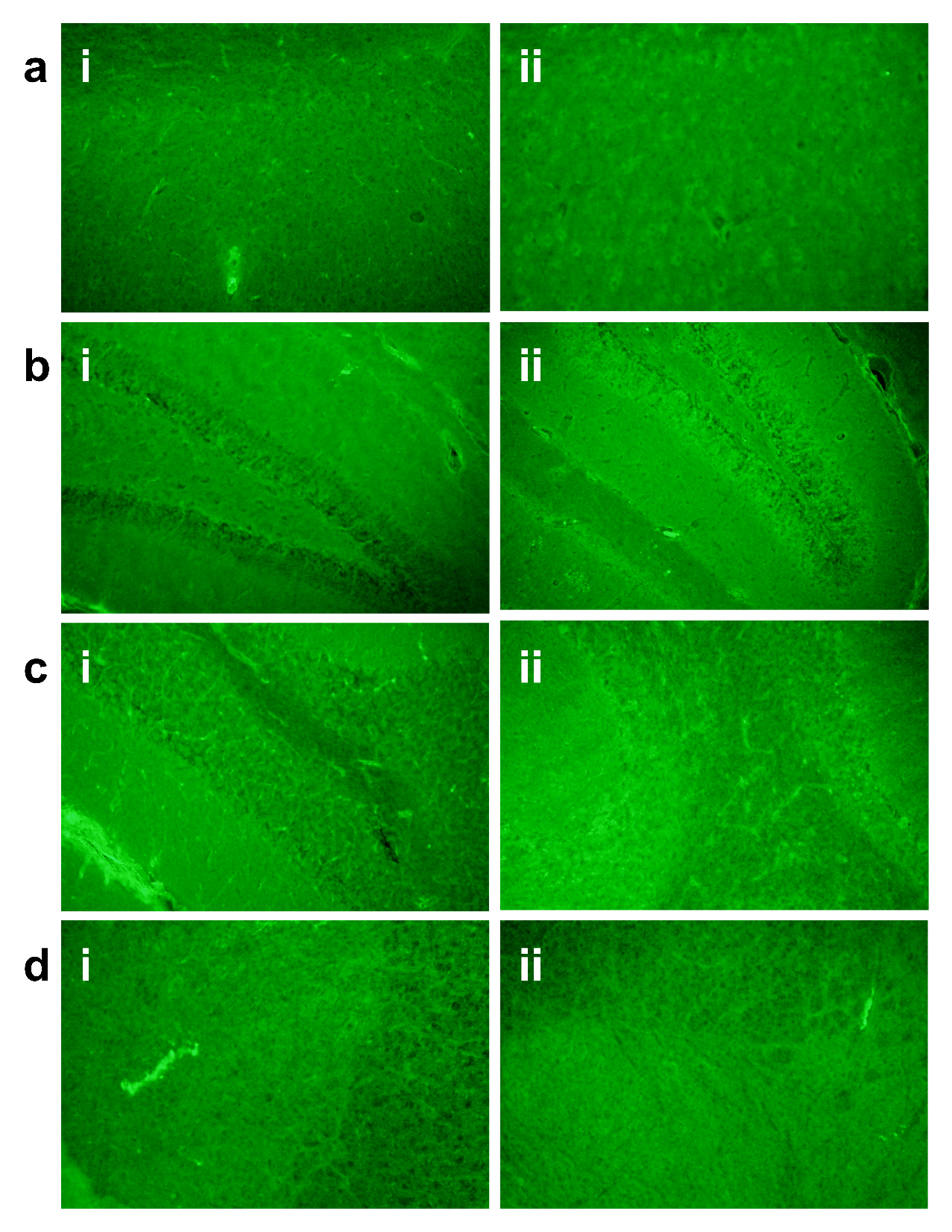

**Figure 3:** Serum albumin labelling in the aging CNS. Example image of serum albumin labelling in the **(a)** cortex, **(b)** hippocampus, **(c)** cerebellum, **(d)** spinal cord in **(i)** young and **(ii)** old. 20x.

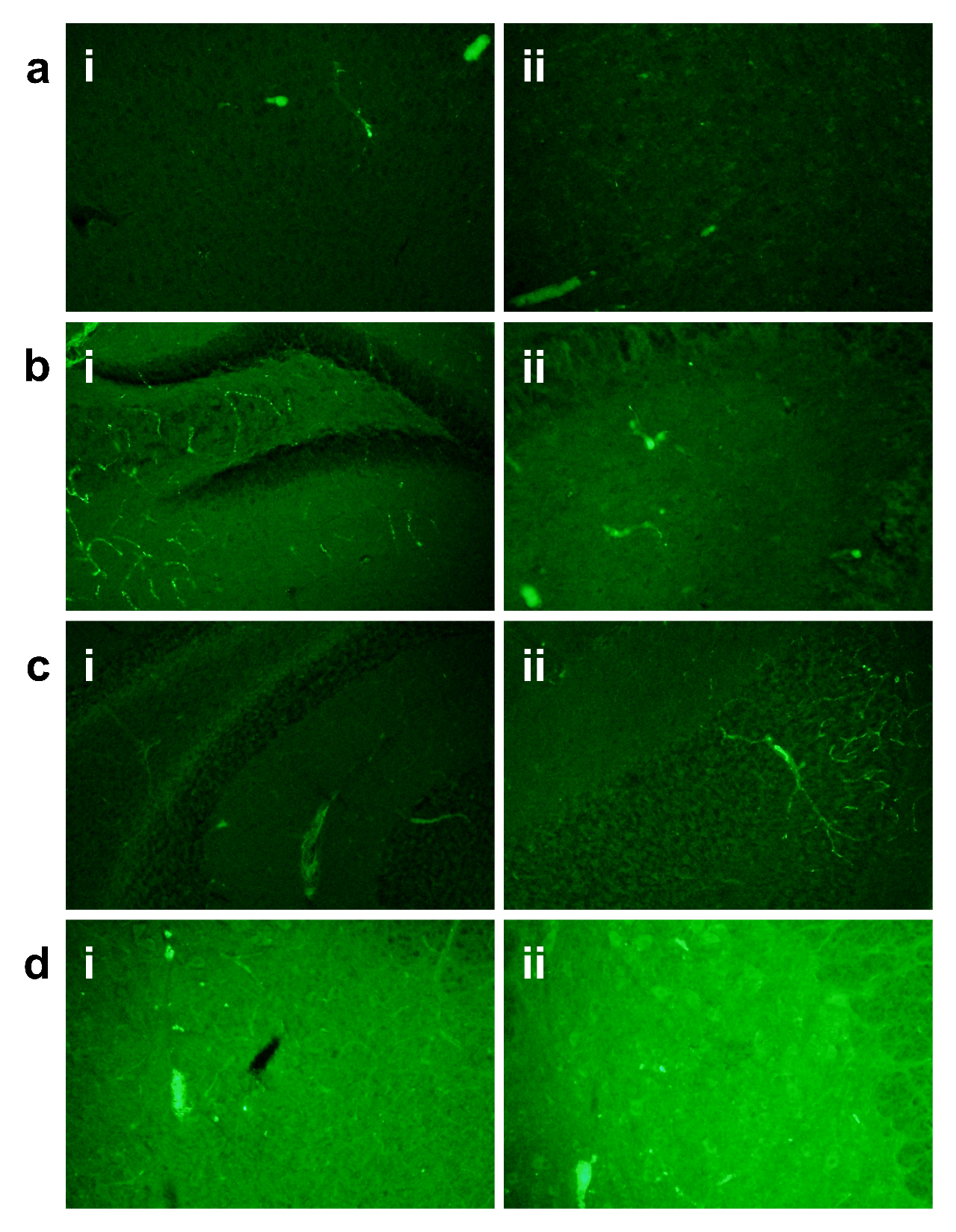

**Figure 4:** IgG labelling in the aging CNS. Example image of IgG labelling in the **(a)** cortex, **(b)** hippocampus, **(c)** cerebellum, **(d)** spinal cord in **(i)** young and **(ii)** old. 20x.

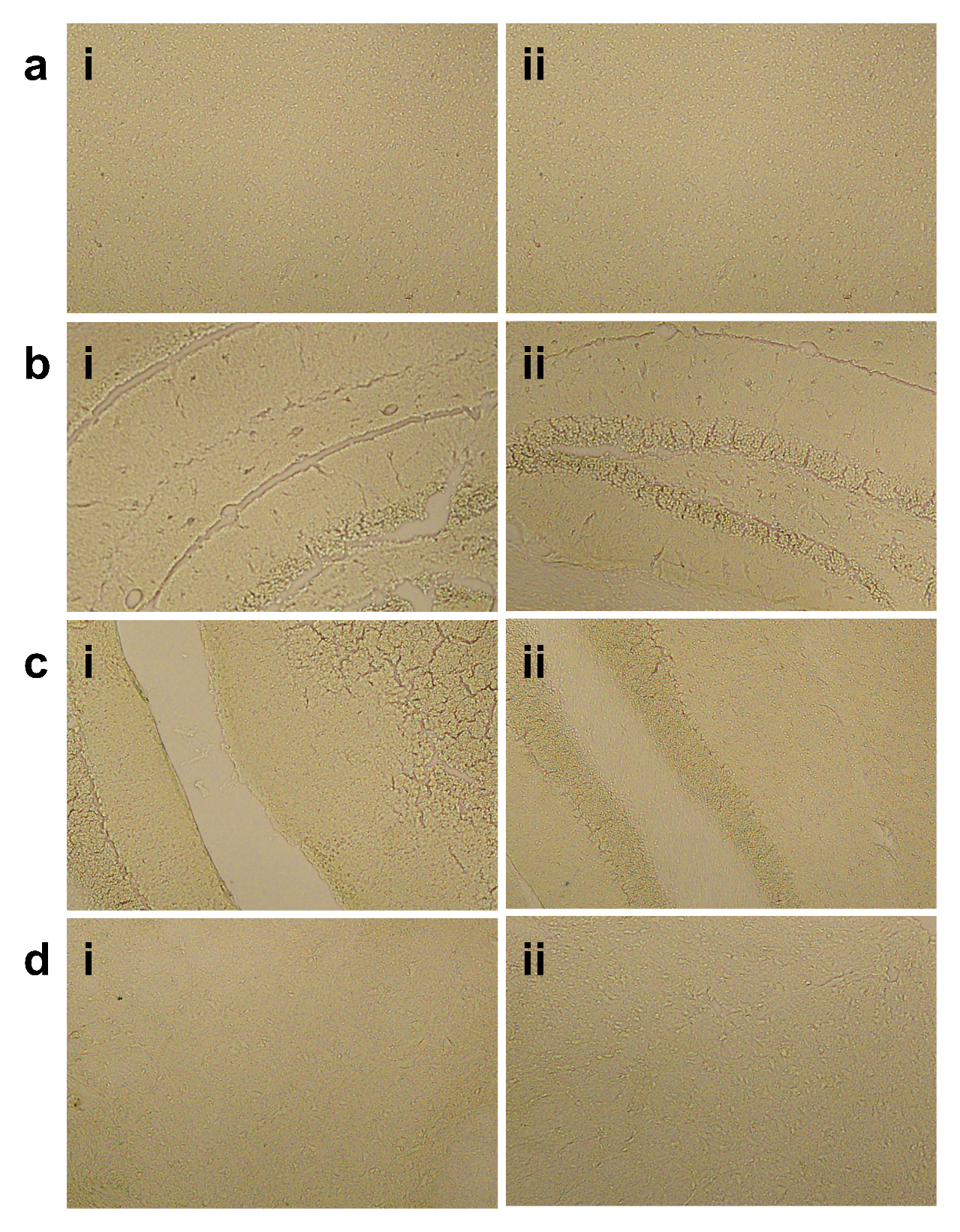

**Figure 5:** Iron labelling in the aging CNS. Example image of iron labelling in the **(a)** cortex, **(b)** hippocampus, **(c)** cerebellum, **(d)** spinal cord in **(i)** young and **(ii)** old. 20x.

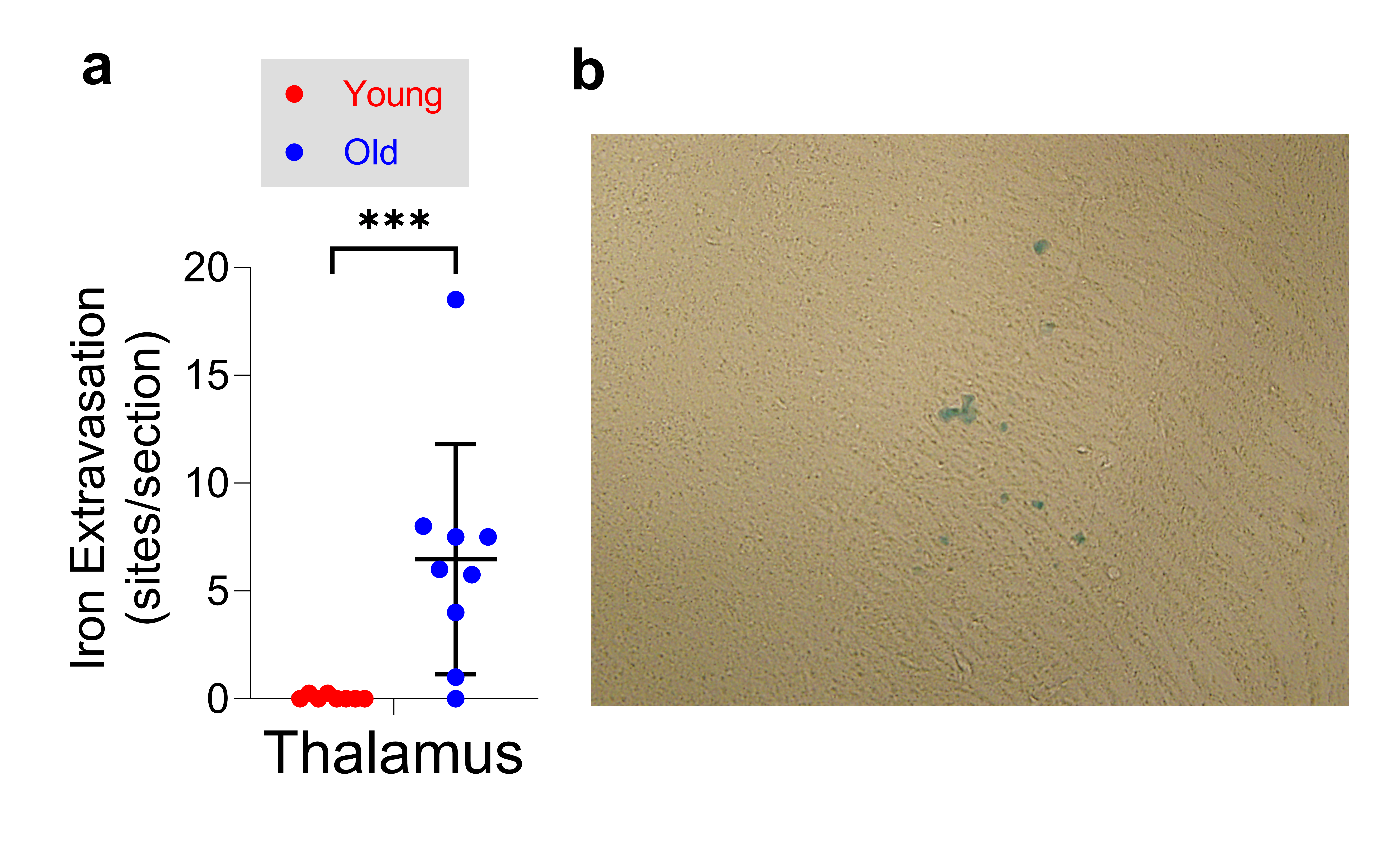

**Figure 6:** Iron labelling in the aging thalamus. **(a)** Number of sites per section of iron extravasation in young and old thalamus (n=8/9/grp). Points are individual animals. *** p-value <0.001. **(b)** Example image of thalamic iron labelling. 10x.

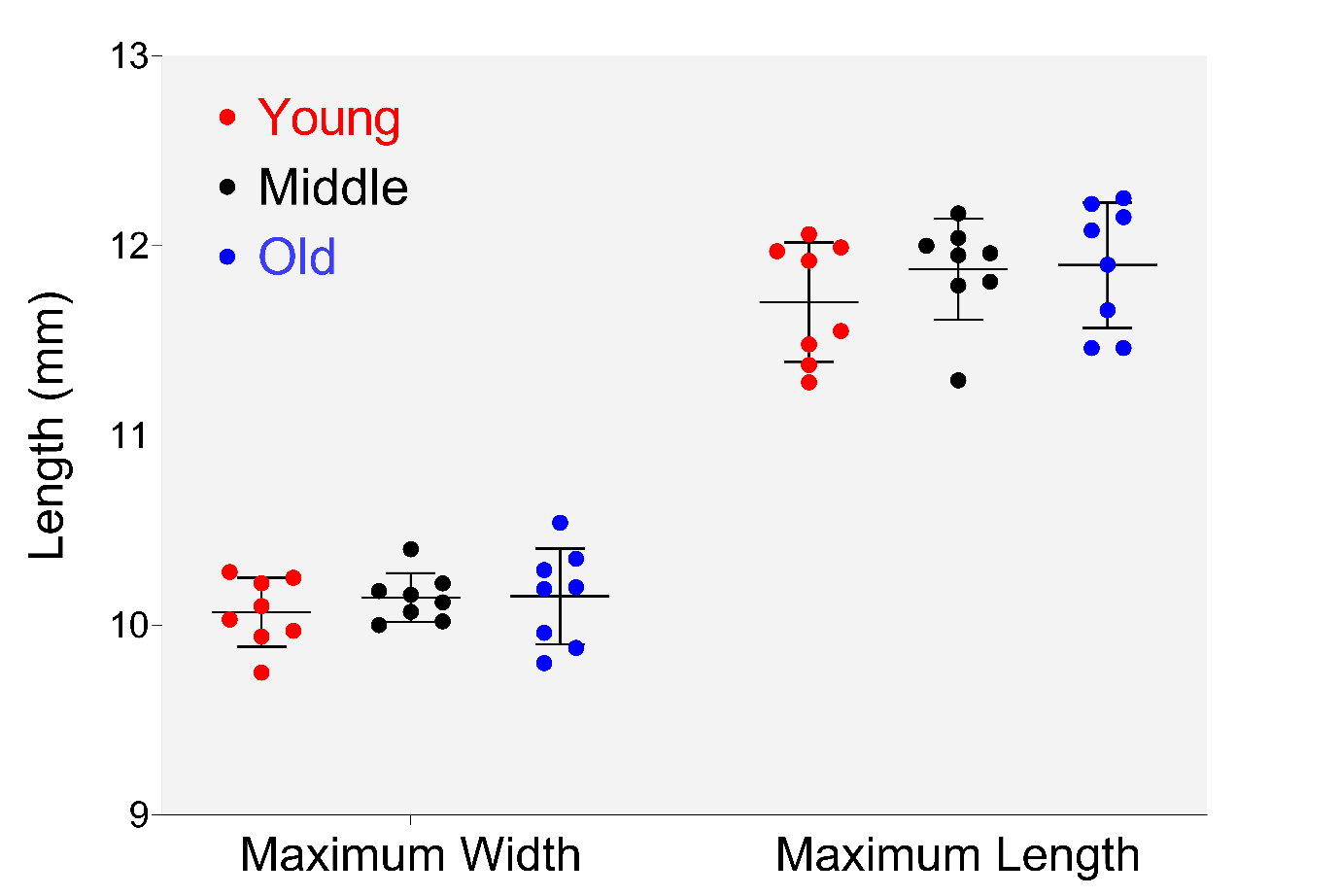

**Figure 7:** Gross measurements of the aging brain. Maximum width and length of the brains from young, middle-age, and old animal (n=8/gp) were measured using calipers. Points are individual animals. No comparisons were significant by Kruskal-Wallis Test.
