## Supplementary File 3 Detailed experimental procedures for "Aging disrupts blood-brain and blood-spinal cord barrier homeostasis, but does not increase paracellular permeability"

Supplementary Experimental Procedures

Antibody Table

| Antibody | Company (Cat Number) | Conjugation | Labelling assay (Dilution) |
| --- | --- | --- | --- |
| Rabbit anti-Col IV | Abcam (ab6586) | Unconjugated | LM (1:100)  Blood vessel density (1:400)  Pericyte coverage (1:1000)  IgG (1:800) |
| Goat anti-CD13 | R&D Systems (AF2335) | Unconjugated | Pericyte coverage (1:500) |
| Rabbit anti-mouse serum albumin | Abcam (ab19196) | Unconjugated | Serum albumin (1:400) |
| Armenian hamster anti-CD31 | Iowa University Hybridoma Bank (2H8) | Unconjugated | Serum albumin (1:400) |
| Donkey anti-rabbit | Jackson Immunoresearch (711-585-152) | Alexa Fluor 594 | LM (1:200)  Blood vessel density, Serum albumin, IgG (1:400)  Pericyte coverage (1:500) |
| Donkey anti-goat | Jackson Immunoresearch (705-545-147) | Alexa Fluor 488 | Pericyte coverage (1:500) |
| Goat anti-Armenian hamster | Abcam (ab175716) | Alexa Fluor 568 | Serum albumin (1:400) |
| Donkey anti-mouse IgG | Jackson Immunoresearch (715-545-151) | Alexa Fluor 488 | IgG (1:400) |

1. Animals

Healthy C57BL/6 male mice were used for all experiments. Mice were maintained under standard housing conditions, on a 12-hour light-dark cycle, with food (Specialty Feeds SF00-100, https://www.specialtyfeeds.com/products/standard-diet/) and water available ad libitum. All mice were euthanized with 1mL i.p. Lethabarb (325mg/ml) before tissue collection. All animal work was undertaken in strict accordance with the University of Newcastle Animal Ethics Committee, and New South Wales and Australian animal research guidelines. Young mice were 2-4 months old (weight range 22-31g, mean±SEM 26.6±0.2g), and old mice were >24 months old (26.5-40g, 32.2±0.3g).

2. Tissue Dissection and Cryosectioning

All phosphate buffered saline (PBS) used was diethyl pyrocarbonate (DEPC) treated and autoclaved before use to inactivate RNases. All perfusions were done with 50mL of each perfusate. For RNA analyses, mice were transcardially perfused on ice with 50mL ice-cold PBS, and brains and spinal cords were dissected out, frozen in isopentane on dry ice and stored at -80°C for later use. Tissues were then mounted in optimum cutting temperature (O.C.T.) compound and cryosectioned at 10µm - 100µm thickness as required. Cryosections were thaw mounted on RNase-free glass microscope slides and stored at -80°C.

For analyses requiring immunolabelling or staining, mice were transcardially perfused on ice with 50mL ice-cold PBS, followed by ice-cold 50mL 4% paraformaldehyde (PFA), pH 7.2. Tissues were dissected out and post-fixed in 4% PFA for 2 hours. Tissues were then cryoprotected in 30% sucrose, mounted in O.C.T. compound, cryosectioned at 40µm, and stored in 0.05% sodium azide in PBS at 4°C for later use.

3. Molecular Analyses – Gene expression

3.1. Bulk RNA Sequencing

a) Tissue preparation for bulk RNA sequencing

RNA sequencing was performed in batches:

Batch 1: spinal cords from 4 young and 4 old mice were obtained as described above. Cervical spinal cords were cut from whole cords and half the cervical region was homogenised and RNA extracted using a Norgen Fatty Tissue kit following manufacturer’s instructions.

Batch 2: Brains from the same animals used for spinal cord analyses, plus one extra brain for each age group, were cryosectioned as described above and frontal cortices (approximately 3 to 2 mm rostral to Bregma) were dissected away from non-cortical tissue, homogenised and RNA extracted as for batch 1.

All RNA samples were DNAse treated in solution using Invitrogen reagents as per manufacturer instructions. Briefly, RNA was DNase treated for 15 mins with DNase I and the DNase subsequently inactivated by addition of 25mM EDTA and heating to 65°C for 10 minutes. DNA-free, RNA samples were sent to the Australian Genome Research Facility (AGRF, Melbourne) for sequencing.

b) Analysis of CNS barrier related gene expression by RNA sequencing

RNA sequencing was carried out to provide lists of differentially expressed genes (DEGs) for analyses for enrichment of barrier-related genes. Total RNA was rRNA depleted, fragmented and first and second cDNA strands synthesised. Adaptors were ligated and the first strand underwent 13 cycles of PCR amplification. cDNA was sequenced using TruSeq PE Cluster Kit v3 reagents and the Illumina HiSeq 2500 system, with between ~82 - 95 million (spinal cord), ~69 - 92 million (cortex) 100 bp, paired end reads, per sample.

Read files were first subject to QC using the FASTQC tool (Andrews, 2010). Adaptors were removed using the Cutadapt tool (Martin, 2011). Forward and reverse reads files were aligned using the STAR aligner (Dobin et al., 2013). Aligned files were assessed to determine DEGs using Cuffdiff (Trapnell et al., 2012). DEG lists were compiled using a FDR cut-off of <0.05. DEG lists were analysed for Gene Ontology (GO) enrichment using the PANTHER Overrepresentation Test for biological process (https://geneontology.org/). DEG lists were compared with the CNS barrier-related gene lists in Supplementary Tables 2-5. Enrichment of CNS barrier-related gene sets in the DEG lists was determined using a hypergeometric overlap calculator: https://systems.crump.ucla.edu/hypergeometric/. p-values were corrected for multiple comparisons.

3.2. Age-related barrier gene expression across CNS regions by qPCR

Brains and spinal cords from 8 young and 8 old mice were cryosectioned at 100µm thickness as described above. Between 6 and 10 slides were taken such that the full rostro-caudal extent of each region of interest was represented. Slides were washed briefly in PBS on ice, dehydrated for 3-5 minutes in 100% EtOH, and the regions to be investigated (cortex; corpus callosum; hippocampus; cerebellum; cervical spinal cord WM; cervical spinal cord GM) dissected out using a sterile scalpel blade under a dissecting microscope. Regions were placed into RNA later (Thermo Fisher) and stored at -80°C for later extraction. RNA was extracted using the miRNeasy Micro Kit (Qiagen) and DNase treated as described above. Reverse transcription (RT) to generate first strand cDNA, was carried out using the Sensifast cDNA synthesis kit (Bioline) according to the manufacturer instructions. Two RT reactions were run, one including the reverse transcriptase enzyme (RT+) and the other without the enzyme (RT-). cDNA samples were diluted to ~0.2 ng/µl in preparation for qPCR analysis.

Genes for qPCR analysis were selected based on importance to CNS barrier function, and on our RNA-seq results. qPCR primers were designed using the NCBI Primer-BLAST program (Ye et al., 2012). Primers were designed across exon-exon junctions, and/or towards the 3' end of the transcript where possible. Primer sequences for all genes are listed in supplementary File 8. All qPCR reactions used the SensiFast Low Rox Sybr Green kit (Bioline). Primer pair specificity was determined by melt curve analysis, and annealing temperatures were adjusted to ensure a single qPCR product for each primer pair. All primer pairs were confirmed to have a single peak melt curve.

qPCR reaction volumes were 12µl: 5µL of sample cDNA + 7µL of master mix. Master mix contained 6µL of Sybr Green, 0.25µL primer stock (final reaction concentration was 210nM for each primer), and 0.75µL nuclease free water. RT+ samples were run in triplicate for each gene. All RT- samples were run in triplicate for primer work up, and then as singles for experimental samples. qPCR reactions were run on Applied Biosystems 7500 or QuantStudio 6 Pro qPCR devices, using the following run parameters:

1.	Initial denaturation and polymerase activation step at 95°C for 10 minutes

2.	40 cycles:

a.	95°C for 15 seconds

b.	64°C for 30 seconds

3.	Melt curve:

a.	95°C for 15 seconds

b.	60°C for 1 minute

c.	Gradient temperature increase from 60°C to 95°C

d.	95°C 30 seconds

e.	60°C 15 seconds

RNA integrity was determined using the 3': 5' ratio for the succinate dehydrogenase (SDH) gene for each sample (Nolan, Hands, & Bustin, 2006). Samples with ratios >4 (2 cycle difference) were removed from analysis. Raw qPCR CT values for each gene were subtracted from the geomean of beta actin (*Actβ*), succinate dehydrogenase (*Sdh*), and glyceraldehyde 3-phosphate dehydrogenase (*Gapdh*), to give a normalised ∆CT for each gene in each sample. Relative expression differences between age groups were determined using the comparative Ct method (Schmittgen & Livak, 2008). The ∆∆CT was transformed to its linear form using: ∆∆CT = 2^^-∆CT^ for statistical analyses and fold change calculations. Statistical significance was determined using one-tailed Wilcoxon Mann-Whiney U Exact test in JMP Pro 14. A p-value cut-off of p<0.05 was used to determine significance. All p-values were adjusted for multiple comparisons using the sequential Holm-Bonferroni procedure (Holm, 1979).

As the spinal cord appeared to have the largest number of barrier related changes by RNA-seq and qPCR, this region was the focus of the time course analysis. Six mice per group, across five age groups (2.5 months, 4 months, 8 months, 14 months, and 26 months), had whole cervical spinal cords dissected, and tissue prepared for gene expression analyses. qPCR reactions were run as described above. Differences between age groups were determined using the Kruskal-Wallis Test followed by a post-hoc Steel-Dwass Test with the 2.5 months old group as the control, with p<0.05 for significance.

3.3. CNS blood vessel-specific gene expression using laser microdissection and qPCR

Six young and 6 old mice had brains and spinal cords removed and cryosectioned at 10µm in preparation for laser microdissection (LM), as described above. Three equally-spaced slides were taken for each forebrain and cervical spinal cord region. Unless otherwise specified, all solutions were ice cold. Slides were briefly rinsed in PBS, fixed for 4 minutes in 100% acetone, rinsed again in PBS, then placed into 2M NaCl PBS for RNA protection (Brown & Smith, 2009). Sections were then labelled using rabbit anti-collagen IV (ColIV) polyclonal primary antibody (Abcam, ab6586), diluted 1:100 in 2M NaCl PBS for labelling of blood vessels. Sections were incubated in primary antibody overnight at 4°C in a humidified chamber. Unbound antibody was removed by rinsing in 2M NaCl PBS. Sections were then incubated for 2 hours at 4°C with Alexa Fluor 594 conjugated donkey anti-rabbit polyclonal secondary antibody (1:200 Jackson Immunoresearch 711-585-152), in 2M NaCl PBS. Unbound secondary antibody was removed by rinsing in 2M NaCl PBS, and slides were placed in 2M NaCl PBS in preparation for LCM.

Immediately prior to LM, slides were briefly rinsed in PBS to remove excess salt, dehydrated in 2 changes of 100% ethanol (10 seconds each), and delipidated in 2 changes of xylene (10 seconds each). Both ethanol and xylene were at room temperature to minimise condensation forming on the slide. LM was carried out using a PALM MicroBeam system (Zeiss) that incorporates an Axiovert 200 M inverted epifluorescent microscope. Blood vessel profiles were collected from GM regions only, using the 40x objective, with isolated vessels laser catapulted into a PCR tube cap containing 30µL RLT buffer (RNeasy Micro Kit Qiagen) and β-mercapto-enthanol solution (1% vol/vol). To reduce RNA degradation, slides were used for a maximum of 2 hours after removal from 2M NaCl PBS. Approximately 900 ColIV positive blood vessel profiles were captured for each CNS region of each animal. Only small blood vessels (<10µm diameter) were captured, to enrich for microvessels. Samples were stored at -80°C.

RNA was extracted, DNase treated, and reverse transcribed as described in above, with minor changes. As RNA yields from LCM material are low, only RT+ cDNA reactions were done. qPCR reactions were run as described in above. Primers used for LM qPCR are listed in Supplementary Table 13. The 3': 5' ratios for the *Sdh* gene for all samples are listed in Supplementary Table 14. Normalisation, fold change calculations, and statistical testing were done as described in above with the exception of normalisation to two genes, *Actβ* and *Sdh*, to allow more genes of interest to be assessed.

3.4 Single cell meta-analysis

Publicly available gene-count data and metadata from all studies was downloaded from the Gene Expression Omnibus. Series Identifiers are as follows: 1. Chen et al., 2020 GSE146395; 2. Zhao et al., 2020 GSE147693; 3. The Tabula Muris Consortium., 2020 GSM4505405; 4. Ximerakis et al., 2019 GSE129788; 5. Li et al., 2021 GSE160991. Data were analyzed in R/4.0.1 using Seurat v3.2.2. Young mice in all studies were 2-3 months old, and old mice were 18-24 months old.

For each study, cells with fewer than 200, and greater than 5,000 expressed genes detected were removed. The Seurat tutorial (https://satijalab.org/seurat/articles/integration_introduction.html) for dataset integration was followed. In brief, each dataset was normalized separately using the NormalizeData function, and variable features were selected using the FindVariableFeatures function with the “vst” selection method selecting 2,000 features. Features that were variable across datasets were selected using the SelectIntegrationFeatures function. Anchors for dataset integration were found using the FindIntegrationAnchors function and datasets were integrated using the IntegrateData function. Data was scaled and centered using ScaleData.

Clustering and visualization using the RunUMAP, FindNeighbours, and FindClusters was performed with defaults except for FindClusters which was run with 0.3 resolution. Cluster marker genes were identified using FindAllMarkers for only positive markers with a minimum logfc.threshold of 0.25. Young and old cells in each cluster for each study were counted using the subset function to select age group (young or old) and study name, and table function to select cell cluster metadata for the cell subset. Clusters were identified by searching for cluster marker genes using 2 databases (https://www.brainrnaseq.org/ and http://mousebrain.org). Differential expression testing between young and old cells in each cluster was performed using FindMarkers with grouping by “Age” and subset by cell cluster, with a minimum logfc.threshold of 0.25.

4. Blood vessel density and pericyte coverage

For blood vessel density, PFA fixed brains and cervical spinal cords from 3 young and 3 old mice were cryosectioned at 40 µm thickness as described in section 2 above. Four equidistant sections were taken from each of the cortex, corpus callosum, hippocampus, and cerebellum, and similarly 6 were taken from the spinal cord for immunolabelling to determine blood vessel density. Sections underwent 3 x 10-minute rinses in PBS, followed by 30 minutes of blocking and permeabilization in 5% normal horse serum (NHS, Sigma-Aldrich) and 0.3% Triton X-100 in PBS. Sections were again washed in PBS and then incubated with primary antibody (1:400 anti-ColIV, Abcam ab6586) over two nights at 4°C on an orbital shaker. Sections were then washed 3 times in PBS and incubated for 2 hrs with AF594 fluorescent secondary antibody (1:400 Jackson Immunoresearch 711-585-152). Finally, sections were washed in PBS, mounted on slides in gelvatol, coverslipped, and imaged on a Nikon D-Eclipse C1 confocal microscope.

Z-stacks of regions were captured using the 40x objective, and the area covered by ColIV was quantified using ImageJ, from a maximum intensity composite after manual thresholding to remove background. In general, 9 Z-stacks were captured per CNS region per animal. However, due to the unusual anatomy of the cerebellum, 6 and 3 Z-stacks for GM and WM regions, respectively, were captured per animal. For each region, the area covered by ColIV blood vessel labelling was determined. Differences between groups were assessed using a nested ANOVA with the animal number variable nested within the age group, with a p<0.05 of the age-effect for significance.

For pericyte coverage, PFA fixed cervical spinal cords from 8 young and 9 old mice were cryosectioned at 40 µm thickness. Six equidistant sections per animal were taken for immunolabelling to determine pericyte coverage. Sections were washed, blocked, and permeabilised as above, and then incubated with primary antibodies (1:500 Goat anti-CD13 R&D Systems AF2335, 1:1000 Rabbit anti-ColIV, Abcam ab6586) over two nights at 4°C on an orbital shaker. Sections were then washed 3 times in PBS and incubated with donkey anti-goat AF488 and donkey anti-rabbit AF594 fluorescent secondary antibodies (1:500 Jackson Immunoresearch 705-545-147 and 711-585-152) as appropriate for each primary antibody host species. Finally, sections were washed, mounted, coverslipped, and imaged as above.

Z-stacks of regions (GM and WM) were captured using the 40x objective, and the relative areas covered by ColIV and CD13 were quantified using ImageJ, and the maximum intensity composites for each label after manual thresholding to remove background. 9 Z-stacks were captured per region per animal. Pericyte coverage was calculated as a percentage ((area of CD13 labelling / area of ColIV labelling) * 100). Differences between groups were assessed using a nested ANOVA with the animal number variable nested within the age group, with a p<0.05 of the age-effect for significance.

5. Functional analyses of blood-CNS barrier permeability

5.1 Analysis of CNS water content

Six young and 6 old mice were euthanized, and the brains and spinal cords immediately extracted. Brains and whole spinal cords were individually weighed in (previously weighed) tubes, to determine tissue wet weight, then placed into an oven at 85°C with lids off. After 7 days, lids were replaced and tubes were allowed to cool to room temperature. Tubes were weighed again to give tissue dry weight. Wet weight, dry weight, and CNS water content percentage ((wet-dry)/wet*100) were calculated and compared using a one-tailed Wilcoxon Mann-Whiney U Exact test in JMP Pro 14, with a corrected p-value of <0.05 for significance. The sequential Holm-Bonferroni procedure was used to correct for multiple comparisons.

5.2 Exogenous tracer-based assessment of blood-CNS barrier permeability

a) Sodium Fluorescein (MW 376.27Da)

Ten young and 11 old mice were injected i.p. with 200µl 6% sodium fluorescein (Sigma-Aldrich 518-47-8) in DEPC PBS. The tracer was allowed to circulate for 15 minutes before mice were euthanized. The thorax was flushed with PBS, and then dried. The right atrium was cut, and blood was collected. The mouse was then transcardially perfused with ice-cold PBS. The brains and spinal cords were removed and briefly rinsed in PBS to remove surface. Tissue samples were taken from frontal cortex, corpus callosum, hippocampus, cerebellum, and cervical spinal cord, weighed on an analytical balance, frozen on dry ice, and stored at -80°C. Blood was allowed to coagulate for 30 minutes, before being centrifuged at 2000g for 10 minutes. The serum was removed and stored at -80°C.

To precipitate proteins in preparation for NaFl fluorometry, samples were treated as follows. One µl of serum was diluted 1:100 in PBS, then 900µl of 20 % tri-chloroacetic acid (TCA) (w/v) was added. Five hundred µl of 20 % TCA was added to CNS tissues, and tissues were homogenised using a probe sonicator (Jomar Life Research). Serum and tissues were incubated for 24 hours at 4°C. Samples were spun at 10,000g for 15 minutes, then 300µL of the supernatant was removed, and vortexed with 150µL of 10M NaOH to make the sample alkaline as NaFl fluorescence is pH dependent (Doughty, 2010; Mota, Carvalho, Ramalho, & Leite, 1991). Samples were briefly spun in a microcentrifuge, and 100µL of each sample was pipetted, in triplicate, into a black-sided, clear bottom, 96 well plate (Corning) for fluorometry. Fluorometry was conducted using a FLUOstar OPTIMA microplate reader (BMG Labtech) with an excitation wavelength of 480-10nm and an emission wavelength of 520nm. Sample fluorescence was normalised to sample serum fluorescence and sample tissue weight, using the equation: NF = TF/(SF*TW)

Where:

NF is normalised fluorescence.

TF is the raw tissue fluorescence.

SF is the serum fluorescence.

TW is the tissue weight in mg.

Group differences were analysed using a one-tailed Wilcoxon Mann-Whiney U Exact test in JMP Pro 14, with a corrected p-value of <0.05 for significance. The sequential Holm-Bonferroni procedure was used for multiple comparisons corrections.

b) Fluorescein Dextran (MW 3kDa)

Three young and 4 old mice were injected i.p. with 400µL 5mg/mL 3kDa lysine fixable fluorescein dextran (ThermoFisher D3306) in sterile PBS. The dye was allowed to circulate for 10 minutes before mice were euthanized. Tissues were dissected out and drop-fixed overnight in 4% PFA. Brains and spinal cords were sectioned, and 8 sections per brain region, and 16 sections for spinal cord regions, per animal, were mounted in gelvatol on slides for assessment on an Olympus BX51 epi-fluorescent microscope. For each animal the number of areas demonstrating extravascular leakage were counted and total counts normalised to the number of sections assessed. Group differences were analysed using a one-tailed Wilcoxon Mann-Whiney U Exact test in JMP Pro 14, with a corrected p-value of <0.05 for significance. The sequential Holm-Bonferroni procedure was used for multiple comparisons corrections.

5.3. Endogenous, blood borne indicators of CNS barrier dysfunction

a). Serum Albumin (MW approx. 69kDa)

Brain and spinal cord sections from 8 young and 9 old mice were washed in PBS, permeabilised with 0.3% triton X-100, and labelled with rabbit anti-mouse serum albumin (1:400, Abcam ab19196) and Armenian hamster anti-CD31 (1:400, Iowa University Hybridoma Bank 2H8) in 0.1% triton PBS at 4°C over two nights on an orbital shaker. Sections were then washed 3 times in PBS and incubated in donkey anti-rabbit IgG AF488 (1:400, Jackson Immunoresearch 711-545-152) and goat anti-Armenian hamster AF568 (1:400, Abcam ab175716) in 0.1% triton PBS for 2 hours at room temperature. Sections were then washed 3 times in PBS, mounted on slides in gelvatol and coverslipped. Sections were assessed on an Olympus BX51 epi-fluorescent microscope, the number of areas demonstrating perivascular serum albumin were counted and normalised to the number of sections assessed. One young animal was excluded from the cortex analysis due to a high number of points of extravasation (>28 times the average of the rest of the group). Group differences were analysed using a one-tailed Wilcoxon Mann-Whiney U Exact test in JMP Pro 14, with a corrected p-value of <0.05 for significance. The sequential Holm-Bonferroni procedure was used for multiple comparisons corrections.

b) IgG (MW approx. 150kDa)

Brain and spinal cord sections from 8 young and 9 old mice were washed in PBS, permeabilized and blocked with 0.3% triton X-100 and 3% bovine serum albumin (BSA), and labelled with rabbit anti-ColIV (1:800, Abcam ab6586) and donkey anti-mouse IgG AF488 (1:400, Jackson Immunoresearch 715-545-151) in 0.1% triton, 1% BSA in PBS at 4°C over two nights on an orbital shaker. Sections were then washed 3 times in PBS and incubated in donkey anti-rabbit IgG AF594 (1:400, Jackson Immunoresearch 711-585-152) in 0.1% triton 1% BSA PBS for 2 hours at room temperature. Sections were then washed 3 times in PBS, mounted on slides in gelvatol, and coverslipped. Sections were assessed on an Olympus BX51 epi-fluorescent microscope, and areas of parenchymal perivascular IgG were counted and normalised to the number of sections assessed. Group differences were analysed using a one-tailed Wilcoxon Mann-Whiney U Exact test in JMP Pro 14, with a corrected p-value of <0.05 for significance. The sequential Holm-Bonferroni procedure was used for multiple comparisons corrections.

c) Iron labelling for microhaemorrhages

Brain and spinal cord sections from 8 young and 9 old mice were washed in PBS, then MilliQ H_2_O, and incubated in 5% Potassium Ferrocyanide in 5% HCl for 30 minutes on an orbital shaker. Sections were washed again in MilliQ H_2_O, then mounted and dried onto gelatin-coated slides, dehydrated through and ethanol gradient (70%, 90%, 100%) and delipidated in 2 changes of xylene. Slides were covered in ultramount, coverslipped, and assessed on an Olympus BX51 microscope under bright field illumination. The number of microhaemorrhages were counted and normalised to the number of sections assessed. Group differences were analysed using a one-tailed Wilcoxon Mann-Whiney U Exact test in JMP Pro 14, with a corrected p-value of <0.05 for significance. The sequential Holm-Bonferroni procedure was used for multiple comparisons corrections.
